# Machine learning enables discovery of DNA-carbon nanotube sensors for serotonin

**DOI:** 10.1101/2021.08.20.457145

**Authors:** Payam Kelich, Sanghwa Jeong, Nicole Navarro, Jaquesta Adams, Xiaoqi Sun, Huanhuan Zhao, Markita P. Landry, Lela Vuković

## Abstract

DNA-wrapped single walled carbon nanotube (SWNT) conjugates have remarkable optical properties leading to their use in biosensing and imaging applications. A critical limitation in the development of DNA-SWNT sensors is the current inability to predict unique DNA sequences that confer a strong analyte-specific optical response to these sensors. Here, near-infrared (nIR) fluorescence response datasets for ~100 DNA-SWNT conjugates, narrowed down by a selective evolution protocol starting from a pool of ~10^10^ unique DNA-SWNT candidates, are used to train machine learning (ML) models to predict new unique DNA sequences with strong optical response to neurotransmitter serotonin. First, classifier models based on convolutional neural networks (CNN) are trained on sequence features to classify DNA ligands as either high response or low response to serotonin. Second, support vector machine (SVM) regression models are trained to predict relative optical response values for DNA sequences. Finally, we demonstrate with validation experiments that integrating the predictions of ensembles of the highest quality CNN classifiers and SVM regression models leads to the best predictions of both high and low response sequences. With our ML approaches, we discovered five new DNA-SWNT sensors with higher fluorescence intensity response to serotonin than obtained previously. Overall, the explored ML approaches introduce an important new tool to predict useful DNA sequences, which can be used for discovery of new DNA-based sensors and nanobiotechnologies.

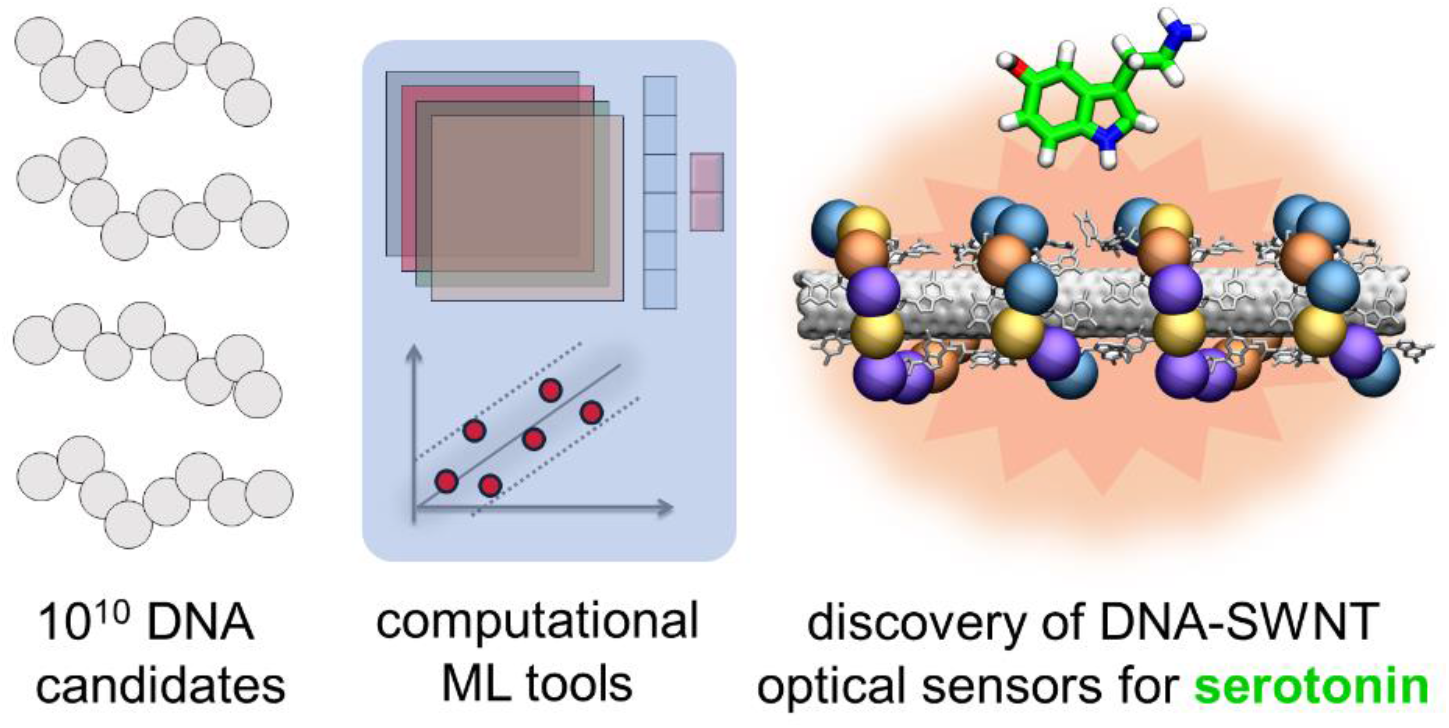

## 1. Introduction

Single walled carbon nanotubes (SWNT) are constituent parts of many new hybrid material systems designed for nanotechnology applications, such as sensing, biological imaging, electronics, and gene delivery^1–10^. Noncovalent polymer adsorption is a widely used method to functionalize SWNTs, while also solubilizing them in aqueous environments by forming a “corona phase” on the SWNT surface. A variety of polymers have been used for SWNT functionalization, including nucleic acids, peptides, surfactants, lipids, and peptoids^11–20^. Among those, nucleic acid functionalized SWNT conjugates are the most ubiquitous and arguably the most technologically useful in important applications, including optical sensing of biologically important analytes^1,2^, polynucleotide (DNA/RNA) delivery for genetic transformation applications^7,21^, and for chirality sorting of multi-chirality SWNT samples into chirality-pure constituents^22–27^.

DNA sequence plays an essential role in DNA-SWNT conjugates that optically sense analytes and is solely responsible for analyte-specific molecular recognition. An effective sequence must simultaneously bind with high affinity to the analyte and the underlying SWNT surface to result in a significant selective change of the SWNT near-infrared (nIR) fluorescence response, ΔF/F, in the presence of the target analyte. Prior work has found that as little as a single nucleotide substitution in a DNA sequence can abolish sensor response to a target analyte^2^.

Most DNA-SWNT-based sensors are generated using either pre-existing molecular recognition elements^28,29^ or low-throughput screening approaches, in which fewer than a hundred DNA sequences are screened for fluorescence modulation by target analytes^2,30^. The latter approach, characterized by random successful parameter hits, relies upon the fortuitous discovery of candidate sensors. While this approach can be useful for starting new directions of research, it is not a sustainable method for optimizing identified sensing technologies or to develop sensors for elusive analytes. In a recent advance, we started addressing this challenge: we developed a method (called SELEC) by which to ‘evolve’ ssDNA-SWNT based molecular recognition towards an analyte of interest, with selectivity that increases with each round of evolution^31^. In this approach, ~10^10^ unique ssDNA strands can be evolved for molecular recognition of target analytes *while still adsorbed to the surface of a nanomaterial*.

The datasets generated by the SELEC approach contain rich information on DNA sequences that confer analyte selectivity and SWNT binding affinity. Herein, we leverage these unique datasets to guide our selection of a new dataset of ~100 DNA-SWNT conjugates for which we determine ΔF/F nIR fluorescence response to the chosen analyte. The resulting dataset is used to develop machine learning (ML) models that learn and make predictions of useful ssDNA sequences - which previously eluded experimental validation - that bind to and optically sense the chosen analyte on SWNT surfaces. The model predictions are examined in validation experiments, the results of which are then used to retrain models and predict new DNA sequences that produce higher ΔF/F response to the target analyte. While our approach could be applied to other analytes, we here demonstrate our approach for serotonin (5-hydroxytryptamine, 5-HT), a neurotransmitter with many roles in the central nervous system^32^ and outside the brain^33^. As serotonin biosensing is of great importance, many recent efforts have been devoted to its sensor development^28,31,34,35^.

## 2. Results

### 2.1. Classifying DNA sequences in DNA-SWNT conjugates based on their optical response to serotonin

We first sought to train and test classifier models for predicting 18-nucleotide (nt) long ssDNA sequences with a high relative nIR fluorescence response to serotonin following conjugation to SWNT. First, models were trained on an initial dataset of 96 unique ssDNA sequences, identified by previous SELEC experiments. This initial dataset was selected from a library of all possible ~10^10^ 18-nt ssDNA sequences that competitively bind to either SWNT (control samples) or SWNT in the presence of serotonin (experimental samples)^31^. The SELEC protocol, schematically shown in **Figure 1a**, was performed for several selection rounds, each of which provided datasets of selected DNA sequences and their abundance. The 96 most abundant sequences from the experimental and control groups from SELEC rounds 3 to 6 were chosen for follow-up serotonin response spectroscopic measurements, thus forming the initial dataset for model training. nIR fluorescence emission was measured for those 96 unique ssDNA-SWNT conjugates before and after the addition of 100 μM serotonin; these data were already reported in Ref.^31^ and are provided in **Figure 1b** and **Table S1**. The response of conjugates to serotonin was calculated from the fluorescence emission spectra for the (8,6) chirality dominant peak (~1195 nm center wavelength) as ΔF/F = (F_a_-F)/F, where F is the fluorescence signal before addition of serotonin, and Fa is the fluorescence signal after the addition of serotonin (**Figure 1a**). The measured ΔF/F values for sequences in the initial dataset range from 0.2 to 1.9.

**Figure 1.**
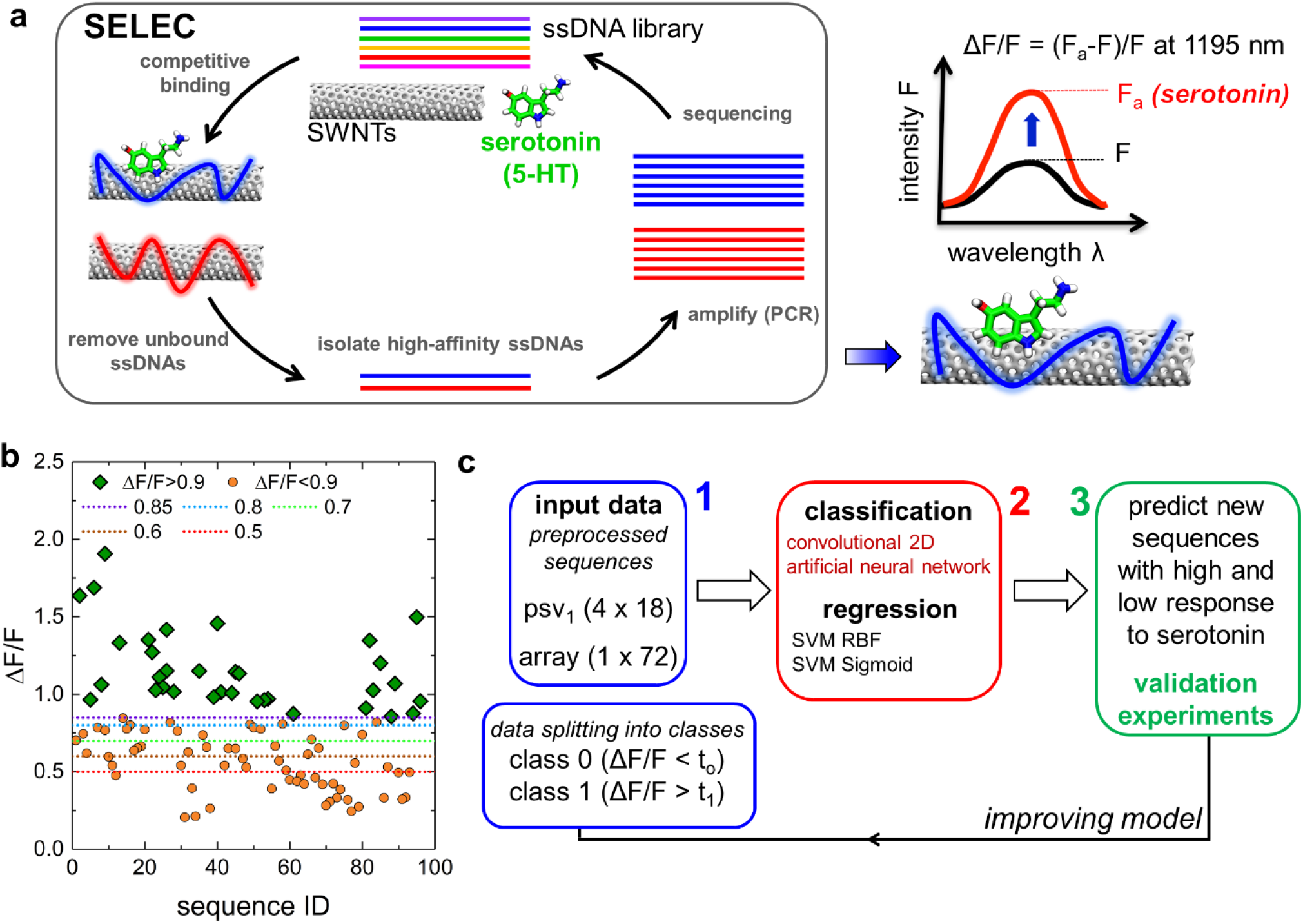
Approach to learning which DNA sequences in DNA-SWNT conjugates provide high response to serotonin. a) A selective evolution protocol, SELEC, performed for up to 6 rounds, experimentally identified ssDNAs with high affinity for SWNTs and, separately, SWNTs in the presence of serotonin. Some of the high-affinity ssDNAs are selected for follow up fluorescence emission spectroscopy experiments of ssDNA-SWNT conjugates before and after the addition of 100 μM of serotonin. b) The optical response, ΔF/F, of 96 unique ssDNA-SWNT conjugates to 100 μM serotonin; the data also include duplicate measurements for 3 of the sequences. c) The main computational approach. DNA sequences are pre-processed into either binary psv1 format or a simple binary array. The sequences are split in two classes of optical response, according to ΔF/F-based threshold values. The sequences and their ΔF/F values are used to train classification and regression models. The models of highest quality are used to predict new sequences with high and low response to serotonin, which are tested in validation experiments. The newly obtained experimental data are then used to generate new models.

Using the approach in **Figure 1c**, convolutional neural network (CNN) classifier models were trained and tested on the obtained dataset of 96 DNA sequences and their corresponding ΔF/F values (**Table S1**, **Figure 1b**). The dataset was split into two classes of sequences based on the response to serotonin, namely, class 1 sequences with a strong response to serotonin (ΔF/F threshold t_1_ > 0.9), and class 0 sequences with a low response to serotonin (variable threshold ΔF/F values of t_0_ < 0.85, 0.8, 0.7, 0.6, and 0.5). Input for CNN models consisted of ssDNA sequences, converted to position specific vector (psv1) form with binary values (example shown in **Figure S1**). The output of trained CNN models are probabilities for the input sequences to belong to class 0 and class 1 (independent).

Quality parameters for one of the best CNN models trained on the initial dataset, model M_1_, are provided in **Figure 2a**. For M_1_, values of the area under the receiver operating curve (AUC) were 0.59 for predicting class 0 sequences, and 0.64 for predicting class 1 sequences, while precision/recall were 0.81/0.5 and 0.76/0.57 for predicting class 0 and class 1 ssDNA sequences, respectively. Models were sensitive to removal of several sequences from input, especially class 1 sequences, as observed when seeking a high-quality CNN model trained on a truncated initial dataset of only 93 datapoints. Quality parameters for a representative model M_1B_, prepared with 93 datapoints from the initial dataset, are reported in **Figure S2**. Overall, we decided to use CNN approach for classifying DNA sequences because several other ML methods tested, including AdaBoost, logistic regression, support vector classification, and random forest, consistently led to class 0 and class 1 probabilities of 0.5 ± 0.2, indicating a poor differentiation between sequences with high and low response to serotonin for models trained with these methods (**Table S2**). Separately, we choose psv_1_ encoding of sequences over other types of encoding reported by others^36^ that we also tested herein. Specifically, we found that term frequency vectors with sequence patterns 1, 2, 3, and 4 nucleotides in length (tfv_1_, tfv_2_, tfv_3_, and tfv_4_), frequently resulted in zero values in confusion matrices for testing sequences, which prevented proper assessment of model quality. Null values in confusion matrices were noted in 90% of models for tfv_1_, 100% of models for tfv_2_ and tfv_3_, and 82% of models for tfv_4_ encoding (**Table S3**).

**Figure 2.**
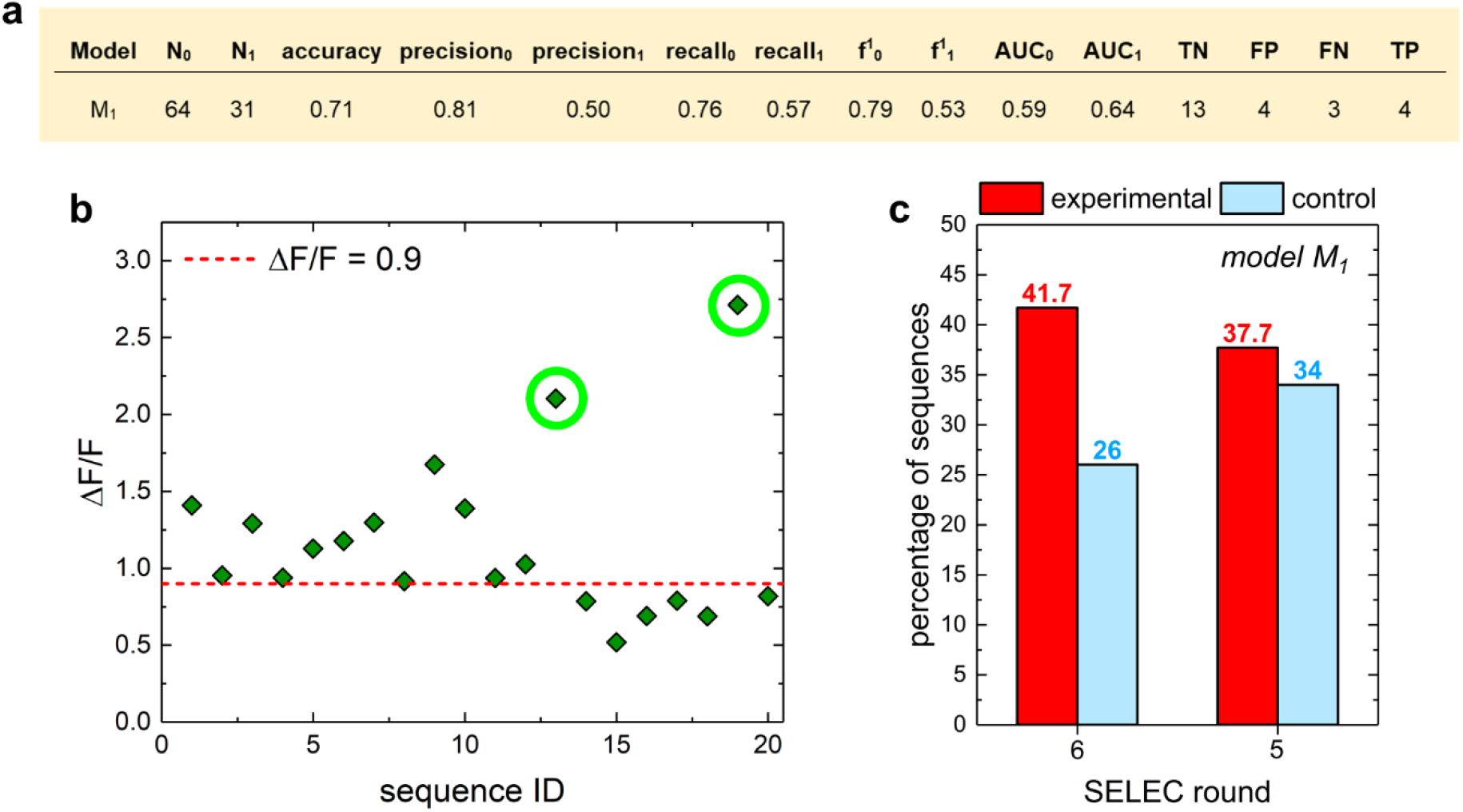
Performance of a representative CNN model trained on the initial dataset. a) Evaluation of a representative CNN M_1_ model trained on the initial dataset, using t_1_ > 0.9 and t_0_ < 0.85. b) The optical response, ΔF/F, of ssDNA-SWNT conjugates to 100 μM serotonin, obtained for 20 new ssDNA sequences, predicted by model M_1_ to have high response to serotonin. Sequences with ΔF/F values exceeding 1.9, the highest values in the initial dataset, are marked with green circles. c) Percentage of sequences in R6E, R5E, R6C, and R5C SELEC datasets predicted by model M_1_ to be high responders to serotonin. The percentage is calculated for the first 300 SELEC data sequences in the given experimental/control dataset that are not in the corresponding control/experimental dataset.

We next examined the predictions of model M_1_ for the most abundant DNAs from control and experimental SELEC datasets. We used model M_1_ to classify the 300 most abundant DNA sequences from round 6 and round 5 experimental (R6E/R5E) and control (R6C/R5C) SELEC datasets, excluding any overlapping sequences found in both experimental and control datasets from the same rounds. Model M_1_ predicts 41.7%/37.7% of R6E/R5E sequences and 26%/34% of R6C/R5C sequences to have high ΔF/F response to serotonin (**Figure 2c**). Since experimental dataset sequences were selected based on their high affinity for SWNTs in the presence of serotonin, in contrast to the control dataset sequences, our expectations were that experimental datasets contain more of the serotonin responsive sequences. These expectations agree with the predictions reported in **Figure 2c**.

To test the quality of model M_1_ predictions, 20 DNA sequences were selected from the 300 most abundant sequences in the R6E SELEC dataset for experimental validation measurements. According to model M_1_ probabilities for these sequences to be in classes 0 or 1, 15 of the sequences are predicted to have high response to serotonin, while the remaining 5 are predicted to have low response to serotonin (**Table S4**). **Figure 2b** and **Table S4** provide the experimentally-measured ΔF/F values for the selected DNA sequences. Interestingly, 12 out of 15 predicted high-response sequences had ΔF/F values greater than the class 1 threshold, t_1_ > 0.9 (80%, obtained from 12 out of 15 sequences). Furthermore, the validation experiments identified two new sequences with ΔF/F values of 2.1 and 2.7, and thus a higher response to serotonin than observed for all the sequences within the initial dataset. These sequences correspond to ID#90 (8 reads, ΔF/F = 2.1) and ID#115 (7 reads, ΔF/F = 2.7), based on the read numbers in the R6E dataset. Separately, 3 out of 5 predicted low-response sequences measured ΔF/F values lower than the class 0 threshold, t_0_ < 0.85. Overall, predicted probability and experimental ΔF/F values do not have a statistically significant correlation (**Figure S3**).

### 2.2. Performance of single CNN models trained on our expanded dataset

To examine if inclusion of additional experimental data points can produce more predictive models, we trained a new representative CNN model M_2_ on the expanded dataset of 113 sequences, which combined the initial dataset (singly measured sequences in **Figure 1b** and **Table S1**) and the additional experimental data from the first set of validation experiments (**Figure 2b** and **Table S4**). A representative model M_2_ has an accuracy of 0.64, AUC values of 0.71 for predicting class 0 sequences, and 0.75 for predicting class 1 sequences, while precision/recall are 0.77/0.59 and 0.53/0.73 for predicting class 0 and class 1 ssDNA sequences, respectively. In addition to improved AUC values, the M_2_ model has significantly improved ROC curves in comparison to models M_1_ and M_1B_ (**Figures 3b, S2**).

**Figure 3.**
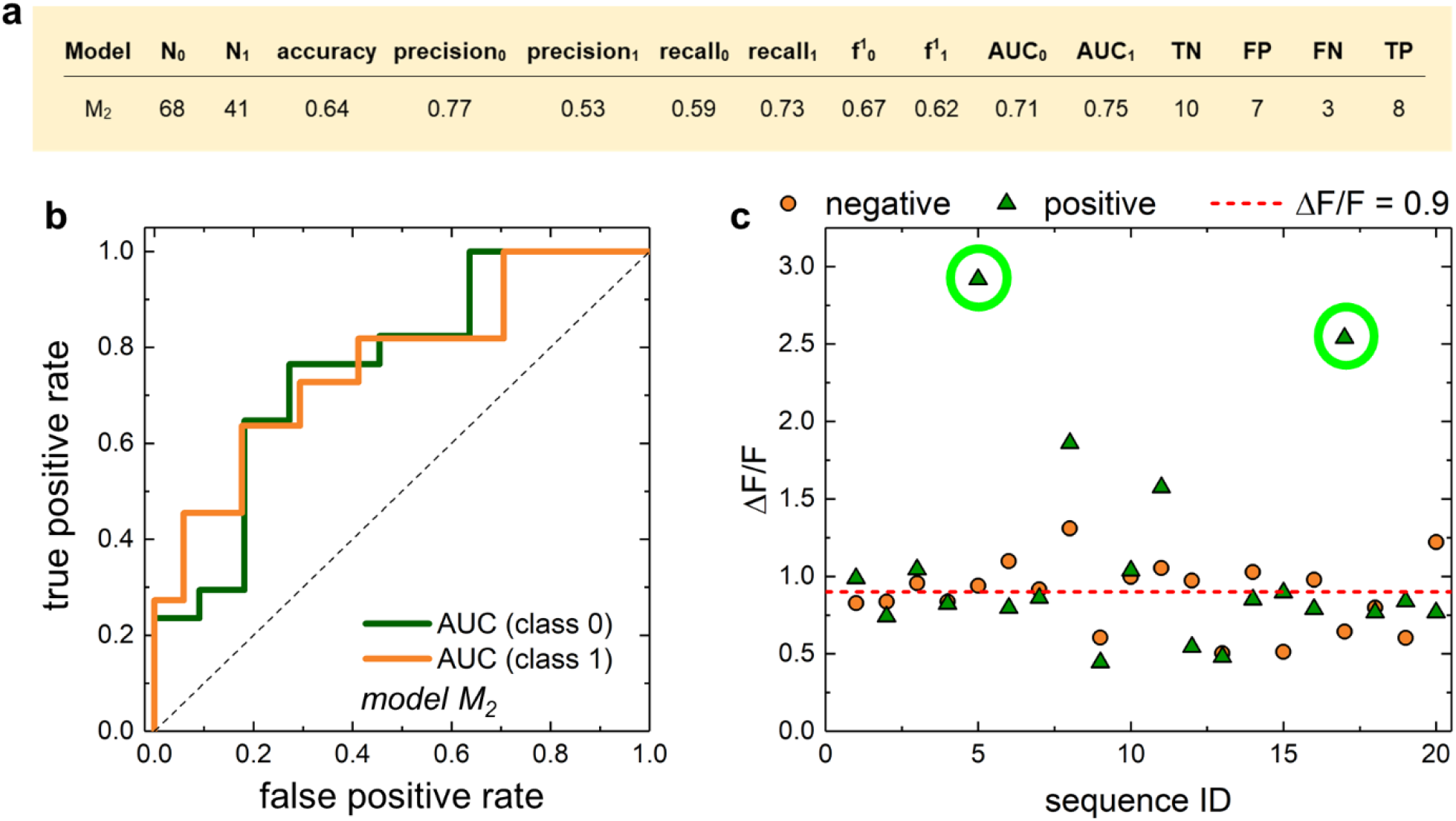
Performance of CNN models using the expanded dataset. a) Evaluation of a representative CNN M_2_ model trained on the expanded dataset, using t_1_ > 0.9 and t_0_ < 0.85. b) ROC curves of model M_2_ when predicting class 0 and class 1 sequences. c) The optical response ΔF/F of ssDNA-SWNT conjugates to 100 μM serotonin, obtained for 40 new ssDNA sequences. Sequences with ΔF/F values exceeding 1.9, the highest value in the initial dataset, are marked with green circles.

To test the quality of model M_2_ predictions, 40 DNA sequences were selected from 280 untested most abundant sequences in the R6E dataset for the next round of experimental validation measurements. Of those 40 DNA sequences, half were predicted by M_2_ to have a low response (labeled as negative) and the other half were predicted to have a high response (labeled as positive) (**Figure S4**). **Figure 3c** and **Table S4** provide the experimentally-measured ΔF/F values for the selected DNA sequences. Model M_2_ overestimates false positive sequences, since only 7 out of 20 sequences (35%) predicted to have high response to serotonin actually have ΔF/F greater than the class 1 threshold of t_1_ = 0.9, while the remaining 13 out of 20 sequences (65%) have ΔF/F < 0.9. Of practical relevance, model M_2_ predicted two previously undiscovered sequences from the R6E evolution group with a very strong response to serotonin, with ΔF/F values of 2.5 and 2.9. Interestingly, these sequences correspond to ID#264 (6 reads, ΔF/F = 2.5) and ID#156 (7 reads, ΔF/F = 2.9), based on the read numbers in the R6E evolution group. Separately, while all the sequences predicted to have a low response to serotonin had ΔF/F < 1.3, only 9 out of 20 sequences (45%) have ΔF/F values smaller than the class 0 threshold of t_0_ = 0.85.

### 2.3. Predicting high response DNAs from combined classification and regression models

When training models M_1_ and M_2_, we noted their stochastic behavior and dependence on the random state variable (different training/testing dataset splits) selected during the training procedure. To characterize the stochasticity of these models trained on our sparse datasets of ~100 sequences, we next analyzed their accuracy and f^1^ scores. The analysis was performed on 200 CNN models trained on the expanded dataset using different random state variables and several t_0_ threshold values (0.5, 0.6, 0.7, 0.8, 0.85). Distributions of these models’ accuracies and f^1^ scores, shown in **Figures 4a-b** and **S5**, range from 0.4 to 0.93 and 0.2 to 0.9, respectively. While these distributions span a wide range, most models have accuracies and f^1^ scores higher than 0.5 and should thus be predictive. Furthermore, more than half of the models in **Figure 4b** have accuracy and f^1^ scores higher than 0.6. Interestingly, the predictions of high response sequences are of higher quality for lower thresholds t_0_ (**Figure S5**, **Table S6**). Model stochasticity decreases and model stability increases once the datasets contain 500 or more sequences per class (**Figure 4c**, input sequences selected from SELEC datasets).

**Figure 4.**
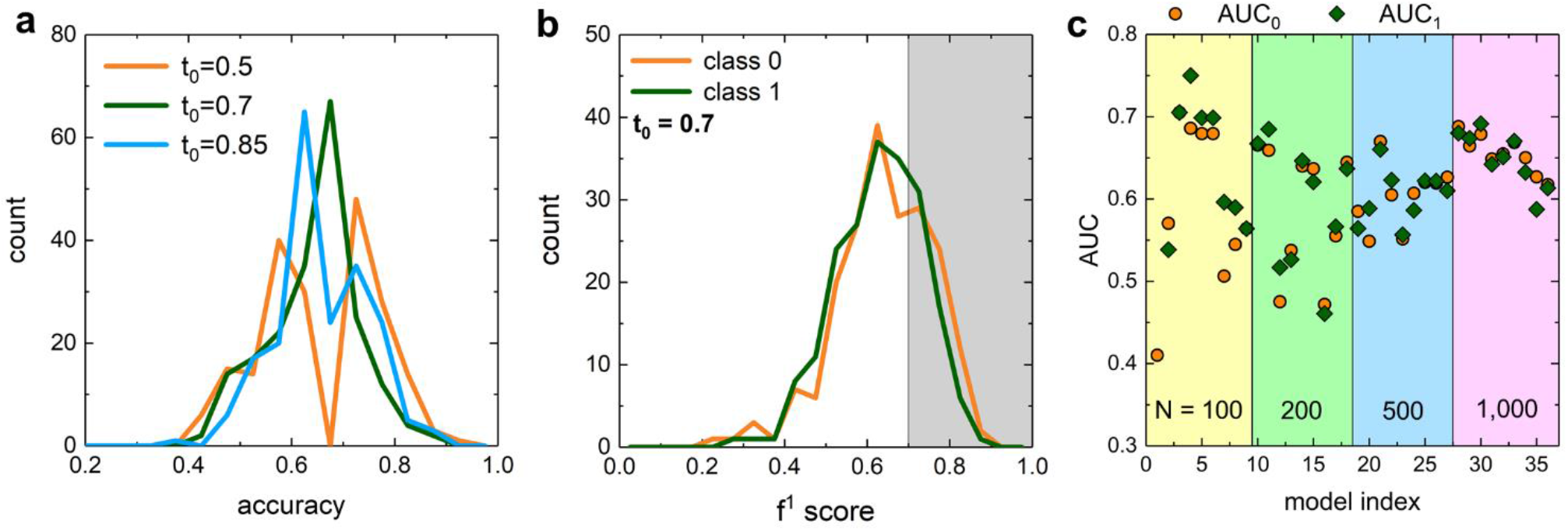
Stochasticity of CNN models. a) Distribution of accuracy values for 200 CNN models with psv_1_ input, obtained using different random states for several t_0_ values. b) Distribution of f^1^ score values for 200 CNN models, obtained using different random states for t_0_ = 0.7. c) Dependence of model stability on dataset size. AUC values for nine CNN models trained on 100, 200, 500 and 1,000 sequences in each of two classes, extracted from R6C (class 0) and R6E (class 1) SELEC datasets. Each model has a different random state variable.

With the objective of predicting DNA sequences with the highest ΔF/F values, we next trained regression models, which predict ΔF/F values based on the DNA sequence input. The regression models were trained using the support vector machine (SVM) regression algorithm with radial basis function (RBF) and sigmoid kernels, based on successful applications of these algorithms for sequence input^37^. One of the best SVM RBF regression models, trained on the expanded dataset with sequences with ΔF/F > 0.9 and ΔF/F < 0.6, is shown in **Figure 5a**. There is a high correlation between ΔF/F values of test sequences obtained experimentally and those predicted by this SVM model, with r^2^ = 0.448 and the Pearson coefficient r_pearson_ = 0.67 (p-value = 0.001). However, as with classification models, the quality of regression models also depends on the random state variable. This dependence is noted in distributions of r^2^ values for 200 models obtained with SVM RBF and SVM sigmoid methods, different random state variables, and different t_0_ values (**Figure 5b-c**). For both SVM RBF and SVM sigmoid methods, r^2^ values range from negative values to 0.5, with SVM RBF models being on average of higher quality than SVM sigmoid models.

**Figure 5.**
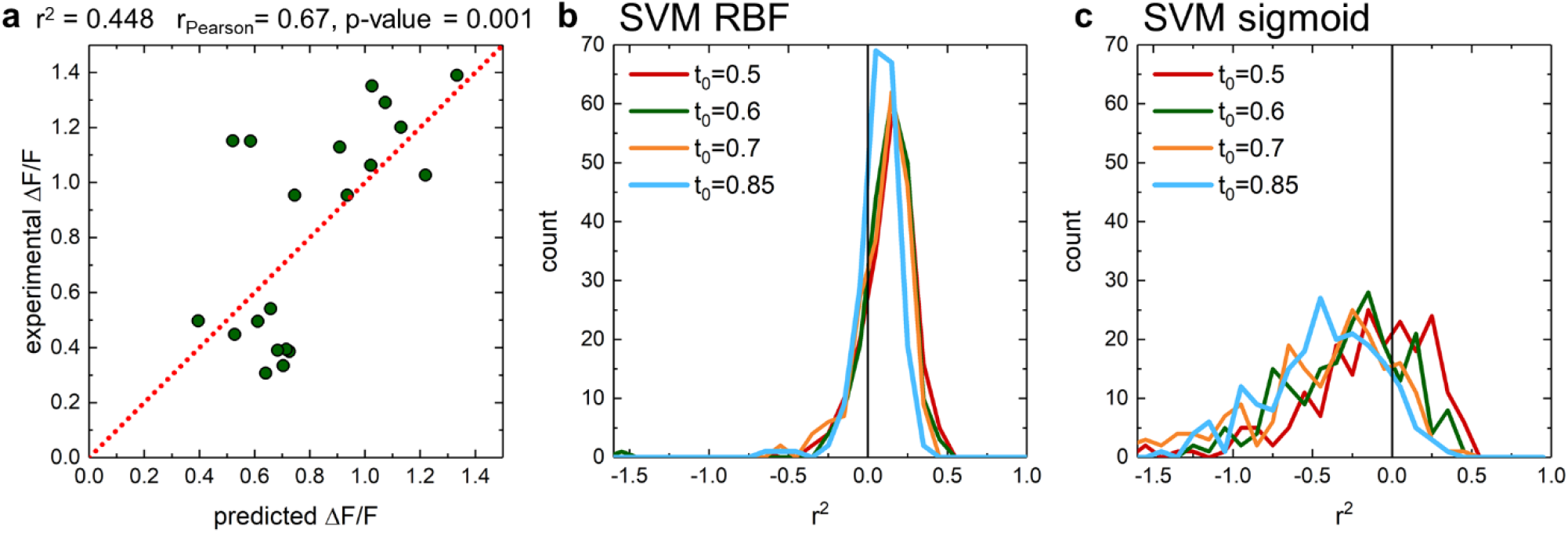
SVM regression models for predicting ΔF/F values of ssDNA-SWNT conjugates. a) Comparison of experimentally measured ΔF/F values and ΔF/F values predicted by one of the best SVM RBF regression models, trained using the expanded dataset and thresholds t_1_ > 0.9 and t_0_ < 0.6 for one selected random state variable. b) Distribution of r^2^ values for 200 SVM RBF models, obtained using different random state variables. c) Distribution of r^2^ values for 200 SVM sigmoid models, obtained using different random state variables.

Given the large number of high-quality classification and regression models obtained, we next assessed whether combining the predictions of these models could be used to determine DNA sequences with high and low response to serotonin. For this purpose, CNN models were trained on input data with thresholds t_1_ = 0.9 and t_0_ = 0.5 or 0.6, which resulted in f^1^_0_ and f^1^_1_ > 0.6. These were next used to predict high and low response sequences from a set of 3,000 most abundant previously untested R6E sequences. Separately, the best regression models with r^2^ > 0.45, trained using the expanded dataset and sequences within thresholds t_1_ > 0.9 and t_0_ < 0.5, were used to predicted ΔF/F values for the same 3,000 sequences. After ranking the sequences according to their regression-predicted ΔF/F values, we extracted the top 10 sequences that are also classified as high response with the CNN models, by having consistently high/low probabilities to be in class 1/class0 (**Figure 6a**). For comparison, we also extracted the bottom 10 ranked sequences, which are also classified as low responders to serotonin based on having consistently low/high probabilities to be in class 1/class0 (**Figure 6a**). The performance of the above top 10 and bottom 10 sequences, labeled as positive and negative, was then examined experimentally (**Figure 6b and Table S7**). 6 out of 10 positive sequences (60%) had ΔF/F response greater than the class 1 threshold of t_1_ = 0.9, and one of them had ΔF/F = 2.1 (**Figure S6**), exceeding the highest value in the initial dataset (1.9). Furthermore, 9 out of 10 negative sequences (90%) had ΔF/F response smaller than the class 1 threshold of t_1_ = 0.9. There is a statistically significant correlation between experimentally measured and predicted ΔF/F values (**Figure 6c**), with Pearson correlation coefficient of r_Pearson_ = 0.5 and p-value of 0.02.

**Figure 6.**
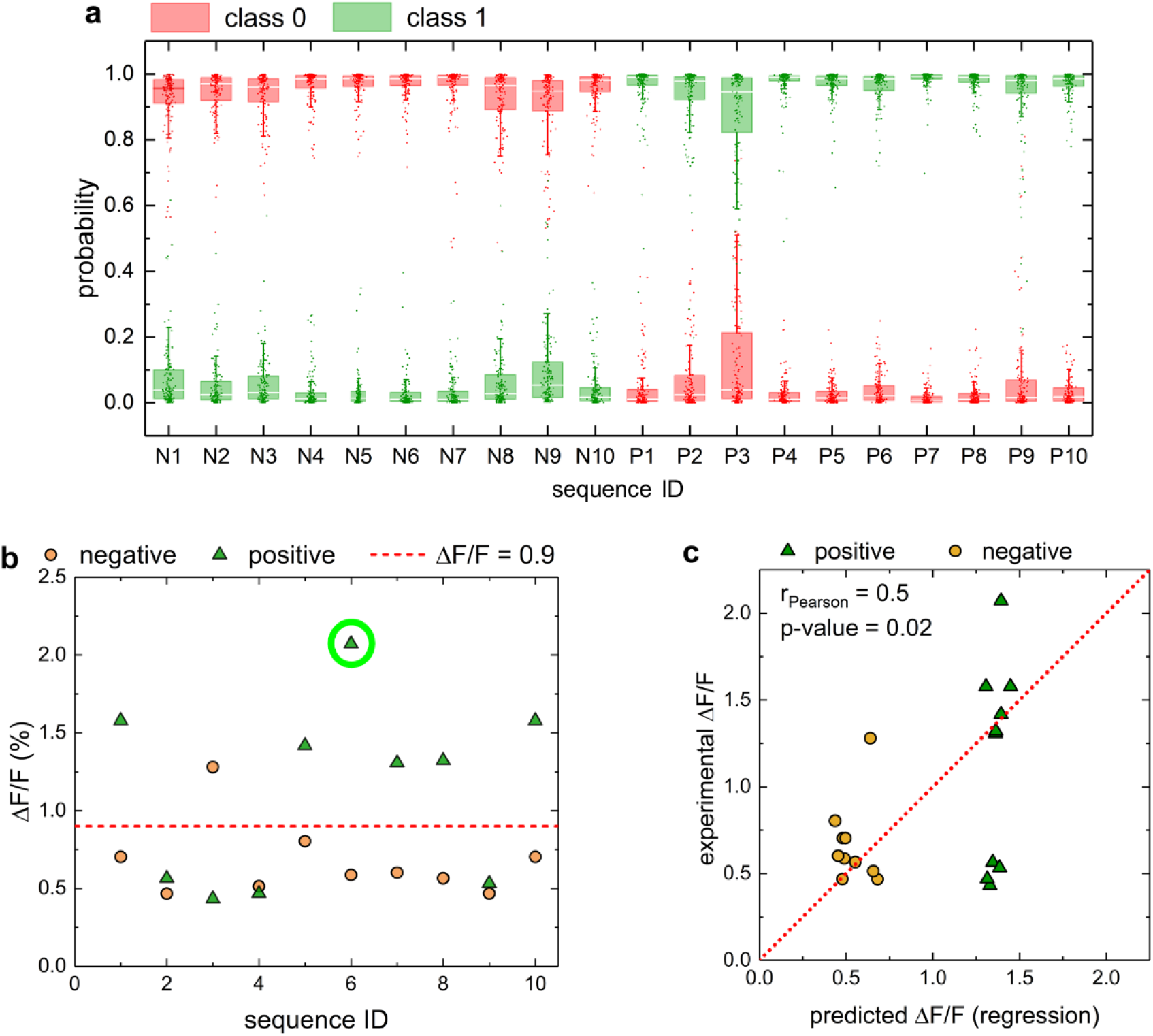
Predicting DNA sequence response to serotonin from multiple high-quality classification and regression models. a) Probabilities of 20 DNA sequences to be high response (class 1) or low response (class 0) to serotonin; these sequences were selected based on predictions of multiple high-quality classification and regression models, as described in the text. b) The optical response, ΔF/F, of ssDNA-SWNT conjugates to 100 μM serotonin, obtained for the same 20 ssDNA sequences. The sequence with ΔF/F value exceeding 1.9, the highest value in the initial dataset, is marked with a green circle. c) Comparison of experimentally measured and predicted ΔF/F values for the same 20 ssDNA sequences.

## 3. Discussion

In this work, we apply machine learning methods towards discovery of new DNA-SWNT sensors of serotonin with high nIR fluorescence response to serotonin. In prior sensor design efforts, we experimentally tested the serotonin response of 96 DNA-SWNT sensors, which were chosen by the abundance (read numbers in library) of each DNA sequence in the experimental and control libraries from SELEC experiments. This previous selection method assumes the best predictor of sensor sensitivity and selectivity is the sequence abundance, which excludes the characterization of low abundance DNA sequences. Here, we demonstrate that ML models can improve on this abundance-based selection. Specifically, ML models can autonomously learn the correlation between DNA sequences and fluorescence responses to analyte and thus assist and improve the selection of better sensor candidates.

ML models can learn the correlations from datasets of already experimentally tested sequences and predict new promising DNA sequences. After testing multiple ML methods, we find that convolutional neural network classifier models provide the most meaningful results when trained on sparse datasets of ~100 DNA sequences, a size typical for new sensor search efforts. At first, we train and test two single CNN classifier models and use them to predict new high response DNA sequences for experimental validation. Even though the prediction accuracies calculated from testing data differ from the accuracies measured in experiments, these models still predicted multiple new DNA sequences with higher response to serotonin than those previously achieved experimentally. Furthermore, we trained and tested regression models on DNA sequence input to predict the relative nIR fluorescence response of these sequences.

Through analyses of model quality parameters for multiple models obtained with different training/testing data splitting, we show that both classification and regression models trained on our sparse datasets are stochastic. However, the majority of models are predictive, since most models have accuracies greater than 50%. Further analyses of model stochasticity dependence on dataset size indicate that model stability can be achieved with datasets of 500 molecules per class. Since obtaining such large datasets in experiments is difficult, we instead explore integrating the predictions of multiple highest quality CNN classifiers and SVM regression models, inspired by model ensembling approaches^38^, and demonstrate an effective increase in the success of this ML approach. We experimentally validate that our integrated approach leads to 60% correct predictions for high responding sequences and 90% correct predictions for low responding sequences, supporting the utility of our method for predicting new promising DNA sequences and accelerating sensor discovery. Separately, we show that a simpler principal component analysis (PCA) approach appears to be predictive in analysis plots but exhibits a poor correlation between predictions and validation experiments (**Figure S7**), in contrast to the successful CNN classifiers and SVM regression models. Furthermore, many sequence patterns are reported in high-response sequences, but most have low abundance (**Table S8**), further confirming the benefit of using ML models when making predictions of high response sequences from existing experimental datasets.

Overall, our ML approaches led to discovery of five new serotonin DNA-SWNT sensors, identified in **Table 1**. Importantly, these sensors all had higher response than sensors previously identified experimentally using only manual screening of the highest-abundance sequences in the R6E SELEC library (ΔF/F = 1.9). Furthermore, the ability to predict DNA sequences that do not respond to serotonin (or any analyte of interest such as interfering agents) with our models is also important for sensor design. Taken together, our results suggest ML approaches can rapidly identify DNA sequences that are great responders for the target analyte and could significantly expedite the development of technologies dependent on DNA-SWNT conjugates, including biosensors, bioelectronics, and chirality separation of SWNTs.

**Table 1.**
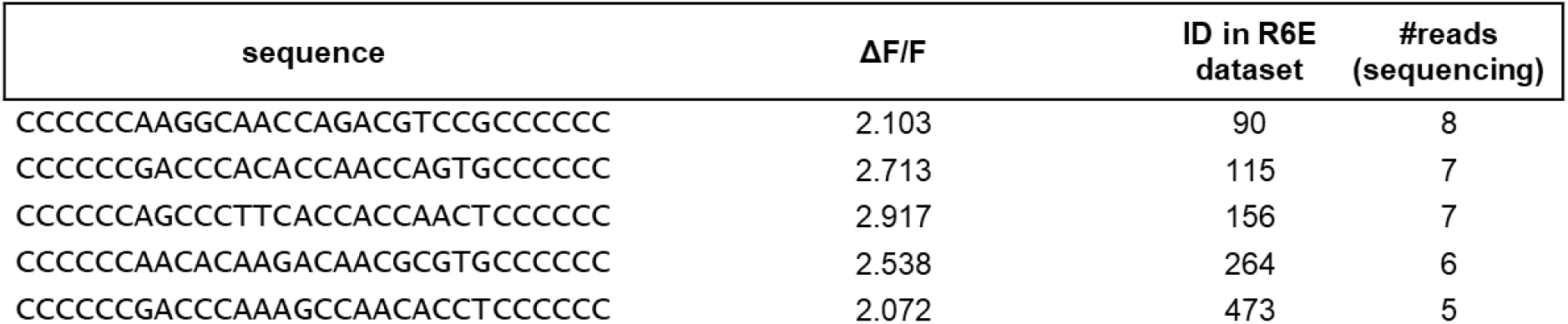
New DNA-SWNT sensors for serotonin. The ID numbers and the numbers of reads are obtained from the R6E SELEC dataset^31^.

## 4. Methods

### Dataset encoding and ML models

Datasets used to train and test the initial ML models consisted of ssDNA sequences and their corresponding ΔF/F values, obtained in experiments reported and described in Ref.^31^ This dataset is also listed in **Table S1**. ssDNAs consisted of 18-nt variable segment flanked by two C_6_-mers from each side. Sequences of 18-nt variable segments of ssDNAs were considered as input data for our models. Several encodings of input data were considered, including position specific vectors (1-gram, labeled as psv_1_ and shown in **Figure S1**) and term frequency vectors (1-, 2-, 3-, or 4-gram, labeled as tfv_1_, tfv_2_, tfv_3_, and tfv_4_ and described in **Table S3**). The dataset from **Table S1** was split into binary classes, where class 0 contains DNA sequences with low response to serotonin and class 1 contains DNA sequences with high response to serotonin. The threshold value to define class 0, t_0_, was varied as a parameter (t_0_ < 0.85, 0.8, 0.7, 0.6, or 0.5). The threshold value to define class 1, t_1_, was held fixed at t_1_ = 0.9. This threshold selection leads to a reasonable balance of class sequences, as required for model training and testing.

Performance of several ML classifier models were tested, including AdaBoost, logistic regression, support vector classification linear, and random forest. For these models, sequences were expressed as 1 × 72 binary arrays, obtained by sequential listing of psv1 matrix columns into a 1-dimensional array. Separately, we tested the performance of convolutional neural network (CNN) models on psv1 and term frequency vector input, successful in previous predictions of DNA and RNA sequence specificities^39^. All our models were trained to predict the probability of the input sequence having high or a low response to serotonin. ML models were trained using scikit-learn library, and CNN models were constructed with Keras and TensorFlow 2 used as backend. ML and CNN models were trained using the initial dataset (**Table S1**). Since the best performance was observed for CNN models with psv1 encoding, the models for extended datasets were generated only with the CNN approach.

All the codes for training ML classification and regression models are freely available on GitHub (https://github.com/vukoviclab/DNAsensor).

### Evaluation Metrics

Our CNN models were trained to predict the response of 18-nt DNA sequences to serotonin, or equivalently, to predict these sequences’ probabilities to belong to class 0 or class 1 molecules. In these models, probabilities for sequences to belong to class 0 and probabilities for sequences to belong to class 1 were evaluated independently. Predicted high response sequences were determined according to the criterion that normalized class 1 probabilities, defined as probability (class1) / [probability (class 0) + probability (class 1)], are greater than 0.5.

For each prepared model, we calculated multiple metrics including accuracy, precision, recall, f^1^ score, ROC curves, and areas under the ROC curves (AUC), and monitored the number of true positives (TP), true negatives (TN), false positives (FP), and false negatives (FN). TP/FP values are numbers of test sequences correctly/incorrectly predicted to have high response to serotonin by the models, and TN/FN values are numbers of test sequences correctly/incorrectly predicted to have low response to serotonin by the models. Accuracy was calculated as Ac = (TP+TN)/(TP+TN+FP+FN), precision was calculated as Prec = TP/(TP+FP), and recall was calculated as R = TP/(TP+FN), and f^1^ score = (2Prec·R)/(Prec+R), according to their standard definitions. For ML models, single values of precision, recall, and f^1^ values were evaluated. For CNN models, two values of precision, recall, and f^1^ scores were reported, allowing the independent assessment of prediction quality for sequences with low and high response to serotonin.

Performance of all the models was also examined with the receiver operating characteristic (ROC) curves, and the areas under ROC curves (AUC). For each CNN model, two ROC curves and AUC values were obtained, one reporting the prediction quality for test sequences with low response to serotonin (AUC_0_), and the other reporting the prediction quality for test sequences with high response to serotonin (AUC_1_).

For most of the datasets (with defined encoding and classification thresholds), 200 models were generated with different random states (different training/testing data splitting). For some sets of the trained models, we report the following evaluations related to model quality metrics: mean, standard deviation, minimum, 25%, 50% and 75% percentile values, and maximum.

### PCA analysis

200 most abundant sequences from R6E and R6C SELEC datasets were analyzed using principal component analysis (PCA) within scikit-learn library. Locations of some of the experimentally tested sequences were then examined in the above determined PCA space.

### Motif search

Sequences from the expanded dataset (used to train and test model M_2_) were split in two classes: sequences that recognize serotonin (positive, ΔF/F > 0.9) and sequences that do not recognize serotonin (negative, ΔF/F < 0.85). These two classes were used to search for DNA sequence motifs associated with serotonin recognition using MERCI software^40^. In the search, minimal occurrence frequency for positive sequences fP and the maximal occurrence frequency for negative sequences f_N_ were set to 3 and zero, and the maximum motif length was set to 18.

### ssDNA-SWCNT suspension preparation

ssDNA-functionalized SWNT suspensions were generated with the following protocol: 1 mg of HiPCo SWNT (NanoIntegris) was added to 0.9 mL of PBS buffer, and the solution was mixed with 100 μL of 1 mM ssDNA. We prepared colloidal suspensions of SWNTs with the initial 96 ssDNA sequences (**Table S1**) and all the subsequent sequences (**Tables S4, S5, S7**) comprising variable 18-nt sequences flanked by two C_6_-mers from each side. The resulting mixture was bath-sonicated for 2 min and tip-sonicated for 10 min at 5-W power in an ice bath. After sonication, the black ssDNA-SWNT suspension was centrifuged for 30 min at 16,100*g* to precipitate nondispersed SWNT, and the supernatant containing solubilized ssDNA-SWNT was collected. The supernatant was spin-filtered with 100-kDa MWCO centrifugal filters at 6,000 rpm for 5 min with DNase-free water to remove unbound ssDNA, and the purified solution at the top of the filter was collected. This spin filtration to remove unbound ssDNA was repeated three times. The ssDNA-SWNT suspension was diluted with PBS buffer and stored at 4°C until use. The concentration of the ssDNA-SWNT suspension was calculated by measuring its absorbance at 632 nm with an extinction coefficient for SWNT of 0.036 (mg/L)^-1^ cm^-1^.

### Fluorescence response measurement of sensors to serotonin

Fluorescence spectra of 99 μL ssDNA-SWNT suspensions (10 mg/L) in PBS were measured before and 10 s after the addition of 1 μL of 10 mM serotonin solution for a final serotonin concentration of 100 μM. We analyzed the fluorescence intensity change of the (8,6) SWNT chirality peak (~1195 nm) in this study. ΔF/F was calculated as ΔF/F = (F_a_ – F)/F based on the baseline fluorescence intensity before analyte addition (F) and the fluorescence intensity 10 s after analyte addition for the (8,6) SWNT chirality (~1195 nm) (F_a_). ΔF/F values for the initial 96 ssDNA sequences, ranging from 0 to 1.9, represent the fluorescence response for the 96 most abundant sequences from the experimental and control groups for SELEC rounds 3 to 6. ΔF/F values of the additional tested sequences are reported in **Tables S4, S5, and S7**.

## Acknowledgments

We acknowledge the support of the NSF CBET-2106587 award (to M.P.L. and

L.V.) and the computer time provided by the Texas Advanced Computing Center (TACC). We also acknowledge support of a Burroughs Wellcome Fund Career Award at the Scientific Interface (CASI) (to M.P.L.), a Dreyfus foundation award (to M.P.L.), a Stanley Fahn PDF Junior Faculty Grant with Award # PF-JFA-1760 (to M.P.L.), a Beckman Foundation Young Investigator Award (to M.P.L.), an NIH MIRA award (to M.P.L.), an NSF CAREER award (to M.P.L), an NSF CBET award (to M.P.L.), an NSF CGEM award (to M.P.L.), a FFAR Young Investigator award (to M.P.L.), a CZI investigator award (to M.P.L), a Sloan Foundation Award (to M.P.L.), a USDA BBT EAGER award (to M.P.L), a USDA NIFA Award (to M.P.L), a Moore Foundation Award (to M.P.L.), a Cisco Research Center grant (to M.P.L), and a DARPA Young Investigator Award (to M.P.L.). M.P.L. is a Chan Zuckerberg Biohub investigator, a Hellen Wills Neuroscience Institute Investigator, and an IGI Investigator. J.A. Acknowledges support from the NSF GRFP.

