## Supplementary Information for "Machine learning enables discovery of DNA-carbon nanotube sensors for serotonin"

**Table S1. The initial dataset of DNA sequences conjugated to SWNT and their ΔF/F response to serotonin.** All sequences are used in training and testing model $M_1$, whereas all sequences except those highlighted in yellow, orange, and green colors are used in training and testing model $M_{1B}$. All the sequences are flanked by two $C_6$-mer PCR primers on each side. The sequences and ΔF/F values were originally reported in Ref.[1]

| Seq ID | Sequence | ΔF/F (1195 nm) | Seq ID | Sequence | ΔF/F (1195 nm) |
|---|---|---|---|---|---|
| 1 | ACACACACAACGACGCGG | 0.70223 | 41 | ACCGCATCGACATGTGCT | 1.01393 |
| 2 | AGCACAACACGGCAACCT | 1.63693 | 42 | ACCGCACGAGCCAGTGTG | 0.54137 |
| 3 | AACACACCACAGACTCTG | 0.74523 | 43 | TCACCACATTCCGCTGTG | 0.65097 |
| 4 | ACACACCATCAGACGCCG | 0.61943 | 44 | ACCGAGAGCAGACGATGT | 1.00867 |
| 5 | AGCAGCACACGACACACT | 0.96503 | 45 | AACACCACACACGGCGCT | 1.1451 |
| 6 | ACGCCAACACATTCCGCT | 1.68843 | 46 | GCAGCGTGACTTGACGTG | 1.13463 |
| 7 | AACACACACAGCCGTCCG | 0.78507 | 47 | AACACGGCCCTCATGTCG | 0.58513 |
| 8 | AACACACACAGACGCACG | 1.0623 | 48 | AGCCGTATGCACACCTCA | 0.52833 |
| 9 | AGCACCAGACAGCACACT | 1.9069 | 49 | ACACACCGTTCATCCGCG | 0.805 |
| 10 | ACCACGATCCTCACTCCG | 0.59733 | 50 | GCTGATCGACGACACGTG | 0.78593 |
| 11 | ACGCACCGACAGCACACT | 0.54127 | 45 | AACACCACACACGGCGCT | 0.64953 |
| 12 | ACACCACACCACACCGAT | 0.47627 | 51 | AGCACACTCCACTCCGCT | 0.95393 |
| 13 | ACGACAACCAACACTGTG | 1.33259 | 52 | GCACACACCAGCCGTCTG | 0.77587 |
| 14 | AGCACACTACACACGGCG | 0.8454 | 53 | AACCACACACCGTCCGCT | 0.9622 |
| 15 | ACACCACCTCACGACGTG | 0.77627 | 54 | ACCACACCATCGACGCGT | 0.97047 |
| 16 | ACACCACCAGACACTGCG | 0.80127 | 55 | AGCCACACGACGCGCTCT | 0.39057 |
| 17 | ACCAACACCAGCCGTGCG | 0.63761 | 56 | ACGGCACACACCATCGCT | 0.66563 |
| 18 | ACACACACCACACGTGCT | 0.65277 | 57 | ACGACACTGCACGACGCG | 0.56955 |
| 19 | ACACAACACCCGACGCGG | 0.66363 | 58 | ACGGCAACTCCCATTCCG | 0.8091 |
| 20 | ACACACACAACGACGCGG | 0.77176 | 59 | ACGACACCACACTGCTCT | 0.50943 |
| 21 | GATCCAACCGCTGCCACA | 1.3514 | 60 | ACACAGCATCATTCCGCT | 0.44783 |
| 22 | ACGACGTACACTCCTCCT | 1.27193 | 61 | GCACCAACCAGCCGTCTG | 0.874 |
| 23 | AACCGCATGTACTCTCCG | 1.02657 | 62 | TCACCACATTCGACGGCG | 0.4382 |
| 24 | AACATGCACAGACGTCCG | 1.11015 | 63 | ACCACAAGTGACTGTCCT | 0.47915 |
| 25 | AACCATGCACAACGCGTG | 1.04517 | 64 | GCCGACATGACTCCTCCT | 0.42083 |
| 26 | ACACAACCTGCTCCTCCT | 1.15197 | 65 | ACACACCAATGACCTGTG | 0.61373 |
| 27 | CCCCCCCCCCCCCCCCCC | 0.81727 | 66 | TACCCACACCACACACTG | 0.70733 |
| 28 | ACGCACAATCCGGCACTT | 1.01673 | 67 | ACTGCACATCGACGCGCG | 0.46227 |
| 29 | ACAGACTGCAGTCATGTG | 0.76213 | 68 | ATTGCCGCCATCCTCATG | 0.6527 |
| 30 | ACACCAGCCACACGTGCG | 0.54147 | 69 | AGGCCACCGTCGCACGTG | 0.41914 |
| 31 | ACCTGACACGATCCTATG | 0.20597 | 70 | ACAGACCGACGTGTGCTG | 0.28237 |
| 32 | GGCACAACGCTCGATGCT | 0.62686 | 71 | TGGGAGCCATCTTGTGCG | 0.30723 |
| 33 | ATTACAGCGGACAAGTGT | 0.39338 | 72 | GTTCAGCCTTTTCGTTCG | 0.42323 |
| 34 | TAAGGCCGATCCCACTAT | 0.21325 | 73 | GGAATCTCCGGCGTCTAT | 0.33207 |
| 35 | TGACTCCATAACAGTGTG | 1.15087 | 74 | TAGCACAGGTCGTCTATT | 0.38593 |
| 36 | GACACCCTGGACCCGTCG | 0.73723 | 75 | GCCAATATAGCCCTTCCG | 0.79868 |
| 37 | TGGCGTACAAACCGTCTG | 0.65847 | 76 | AATCACTGCAATGGTCGT | 0.31937 |
| 38 | ACACACTCTACTCTTCCA | 0.26397 | 77 | AACACATTGACGTGCACT | 0.2447 |
| 39 | GACGTTGTGCCCAAGTTG | 0.98223 | 78 | GGGCTGTGCCGTCATGCG | 0.55663 |
| 40 | AAGGGACTGAAAGCAATG | 1.45753 | 79 | GATGGGGAATCATGCGTG | 0.27437 |

| Seq ID | Sequence | ΔF/F (1195 nm) |
|---|---|---|
| 80 | GCACAATCCAGCGCACAA | 0.73998 |
| 81 | ACGACGGAACTACACACC | 0.91085 |
| 82 | GAGACTCAACCGAACACC | 1.3473 |
| 83 | ACCACACAACCGACTGTG | 1.02557 |
| 84 | AACCCCAACCACGGTTGG | 0.82087 |
| 85 | AGGACAACCCCGCGTGTG | 1.2009 |
| 86 | ACACACCGACACGGTGTG | 0.3311 |
| 87 | ACCACGACGACGACTGTG | 0.53127 |
| 88 | ACCAACACACACTCCGCT | 0.85663 |
| 89 | AACACACCAACACCCGCT | 1.0687 |
| 90 | ACACACACACACTCCGCT | 0.49517 |
| 91 | ACACCACCACACTCCGCT | 0.32298 |
| 92 | ACCACACAACGCTCCGCT | 0.334 |
| 93 | ACACACCGCTCTCCCTCT | 0.49743 |
| 94 | ACGACATGGCACACCGAT | 0.87663 |
| 26 | ACACAACCTGCTCCTCCT | 1.41713 |
| 95 | ACACCAATCGCACTTCCG | 1.4976 |
| 96 | ACACGATCCAACACTCCG | 0.95543 |
| 9 | AGCACCAGACAGCACACT | 0.767 |

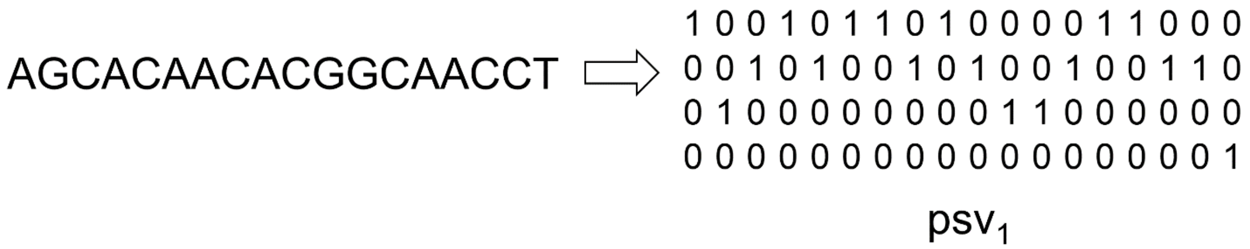


**Figure S1. Example of position specific vector encoding.** The encoding, $psv_1$, shown for an 18-nt DNA sequence is used for training convolutional neural network models.

**a**

| Model | $N_0$ | $N_1$ | accuracy | $precision_0$ | $precision_1$ | $recall_0$ | $recall_1$ | $f^1_0$ | $f^1_1$ | $AUC_0$ | $AUC_1$ | TN | FP | FN | TP |
|---|---|---|---|---|---|---|---|---|---|---|---|---|---|---|---|
| $M_{1B}$ | 62 | 27 | 0.70 | 0.85 | 0.50 | 0.69 | 0.71 | 0.76 | 0.59 | 0.78 | 0.71 | 11 | 5 | 2 | 5 |

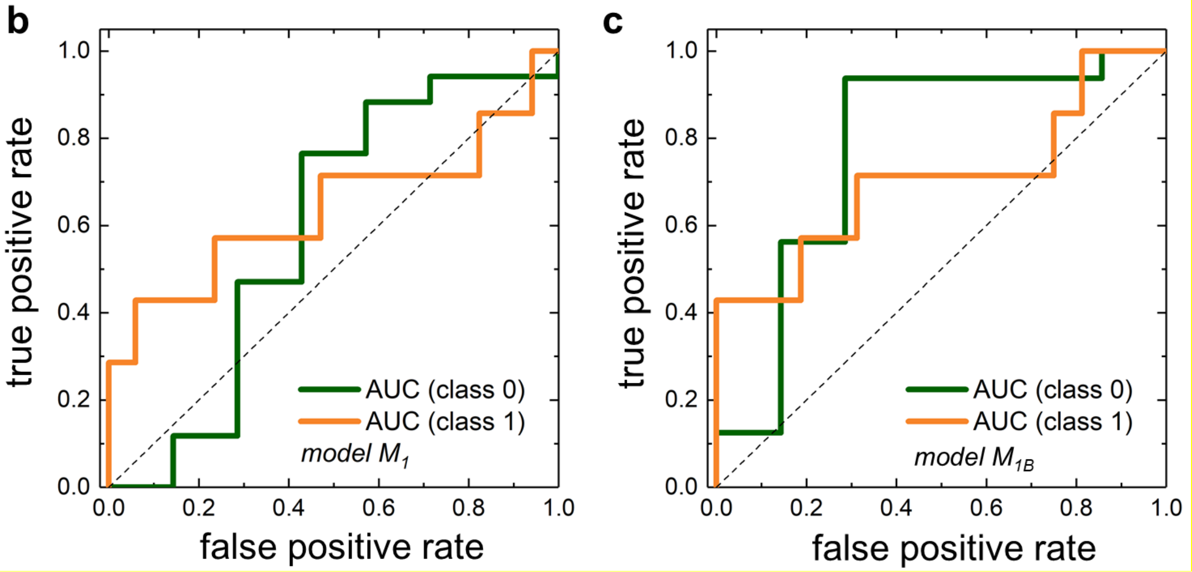


**Figure S2**. **Performance of two representative CNN models trained on the initial dataset.** (a) Evaluation of a representative CNN $M_{1B}$ model trained on 93 unhighlighted sequences from the initial dataset (Table S1), using $t_1 > 0.9$ and $t_0 < 0.85$. (b-c) Receiver operating characteristic curves for models $M_1$ and $M_{1B}$ when predicting class 0 and class 1 sequences.

**Table S2. Example predictions for several ML algorithms.** ML model predictions provide the probabilities for five random sequences to be in class 0 (low response to serotonin) or class 1 (high response to serotonin). The sequences were encoded in $psv_1$ format, and classification of the input data was performed with thresholds $t_1>0.9$ and $t_0<0.85$. While the predictions in this table are shown for only five sequences, the probabilities were in similar range (0.5 ± 0.2) for hundreds of sequences examined.

| | AdaBoost | | Logistic regression | | Support vector classification (linear) | | Random forest | |
|---|---|---|---|---|---|---|---|---|
| **sequence** | **probability (class 0)** | **probability (class 1)** | **probability (class 0)** | **probability (class 1)** | **probability (class 0)** | **probability (class 1)** | **probability (class 0)** | **probability (class 1)** |
| AGCACAGCACGACGCGTA | 0.4867 | 0.5133 | 0.4123 | 0.5877 | 0.6028 | 0.3972 | 0.4741 | 0.5259 |
| AACACCGCACCATCCGAT | 0.4807 | 0.5193 | 0.3794 | 0.6206 | 0.5972 | 0.4028 | 0.3624 | 0.6376 |
| AACGACACACATTCCGCT | 0.4844 | 0.5156 | 0.4920 | 0.5080 | 0.5963 | 0.4037 | 0.5431 | 0.4569 |
| AACCCGACACCACACCTG | 0.4812 | 0.5188 | 0.4392 | 0.5608 | 0.5749 | 0.4251 | 0.6167 | 0.3833 |
| ACACACACCACGCGCGCT | 0.5190 | 0.4810 | 0.6920 | 0.3080 | 0.6247 | 0.3753 | 0.6851 | 0.3149 |

**Table S3. Performance of CNN models with term frequency vector encoding of sequences.** Number of models with non-zero values in confusion matrices out of 200 evaluated models, obtained using term frequency vector (tfv) encoding. Classification of the input data was performed with thresholds $t_1$>0.9 and $t_0$<0.85. Term frequency vector is an array of integer values that count the number of times that segments with uniquely defined sequences occurred within an input 18-nt long sequence. Subscripts 1, 2, 3, and 4 in tfv labels refer to the numbers of nucleotides in these uniquely defined segments. 1, 2, 3, and 4 nt-long segments are defined to cover all possible combinations of A, T, C, and G nucleotides.

| encoding | models with non-zero values in confusion matrix (out of 200) |
|---|---|
| $tfv_1$ | 19 |
| $tfv_2$ | 0 |
| $tfv_3$ | 0 |
| $tfv_4$ | 36 |

**Table S4. DNA sequences, their probabilities predicted by model $M_1$ to belong to either class 0 or class 1, and their ΔF/F response to serotonin.** The sequences predicted to be in class 1 are highlighted in green. The sequences that are experimentally validated to belong to class 1 are highlighted in blue. Normalized probability is defined as probability (class1) / [probability (class 0) + probability (class 1)]. All the sequences are flanked by two $C_6$-mer PCR primers on each side.

| Seq ID | Sequence | Probability (class 0) | Probability (class 1) | Normalized probability (class 1) | ΔF/F (1195 nm) |
|---|---|---|---|---|---|
| M1-P1 | AGCACAGCACGACGCGTA | 0.1094 | 0.8531 | 0.8863 | 1.4098361 |
| M1-P2 | AACACCGCACCATCCGAT | 0.0321 | 0.8876 | 0.9651 | 0.9535565 |
| M1-P3 | AACGACACACATTCCGCT | 0.2369 | 0.7195 | 0.7523 | 1.291181 |
| M1-P4 | AACCCGACACCACACCTG | 0.1554 | 0.5719 | 0.7863 | 0.9386187 |
| M1-P5 | ACCACATCCACAGCCGAT | 0.0252 | 0.8675 | 0.9718 | 1.1286216 |
| M1-N1 | GACCACTCCAATTCCGCT | 0.7245 | 0.5431 | 0.4284 | 1.1771082 |
| M1-P6 | AGCACCACCAGACTCCTG | 0.4073 | 0.7896 | 0.6597 | 1.2978858 |
| M1-P7 | ACGCCACACCATTCCGCT | 0.3686 | 0.6482 | 0.6375 | 0.9159415 |
| M1-N2 | AACCCGAAGCCTGGACCT | 0.5800 | 0.5376 | 0.4811 | 1.6746432 |
| M1-P8 | AGCACAACACGGCACCGT | 0.0447 | 0.8132 | 0.9479 | 1.3900039 |
| M1-P9 | ACAACAACACCTTCCGCT | 0.0280 | 0.8507 | 0.9682 | 0.9373578 |
| M1-P10 | AACACAACAGCTCCTCCT | 0.0179 | 0.9821 | 0.9821 | 1.0271291 |
| M1-P11 | AAGGCAACCAGACGTCCG | 0.0682 | 0.9073 | 0.9301 | 2.1030996 |
| M1-P12 | ACAACCACCGATCCATCG | 0.3599 | 0.5156 | 0.5890 | 0.7858219 |
| M1-N3 | ACACTCTCCCATTCCGCT | 0.5827 | 0.3898 | 0.4009 | 0.5199048 |
| M1-P13 | AACACCACTCGACGCGTA | 0.2072 | 0.6818 | 0.7669 | 0.690589 |
| M1-P14 | AGCCAACATCATTCCGCT | 0.4969 | 0.7996 | 0.6168 | 0.789971 |
| M1-N4 | ACACCCACTCCACACGCT | 0.2909 | 0.2677 | 0.4792 | 0.688906 |
| M1-P15 | GACCCACACCAACCAGTG | 0.3586 | 0.7685 | 0.6818 | 2.713169 |
| M1-N5 | AGCACCCTCTACAGCACA | 0.7011 | 0.5371 | 0.4338 | 0.8194765 |

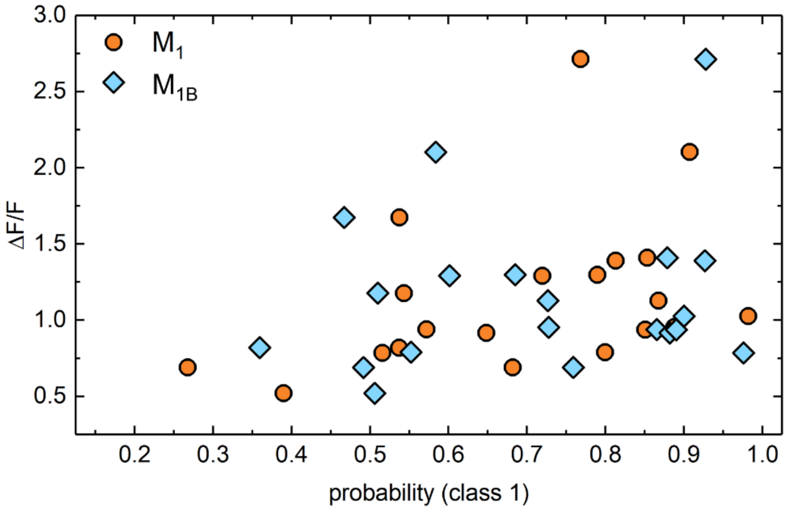


**Figure S3. Correlation between the experimental response and the predicted probabilities of sequences to be high responders to serotonin.** Probability of 20 sequences listed in Table S4 to be in class 1, as determined by models $M_1$ and $M_{1B}$, versus the experimentally measured ΔF/F. Data for models $M_1$ and $M_{1B}$ have Pearson correlation coefficients of 0.383 (p-value 0.096) and 0.164 (p-value 0.49).

**Table S5. DNA sequences, their probabilities predicted by model $M_2$ to belong to either class 0 or class 1, and their ΔF/F response to serotonin.** The sequences predicted to be in class 1 are highlighted in green. The sequences that are experimentally validated to belong to class 0 (top table) or 1 (bottom table) are highlighted in orange and blue, respectively. All the sequences are flanked by two $C_6$-mer PCR primers on each side.

| Seq ID | Sequence | Probability (class 0) | Probability (class 1) | Normalized probability | ΔF/F (1195 nm) |
|---|---|---|---|---|---|
| M2-N1 | ACACATCAACTCCGCTCG | 1.0000 | 0.0000 | 0.0000 | 0.82786 |
| M2-N2 | ACACACCGATCCTACTCG | 1.0000 | 0.0000 | 0.0000 | 0.83695 |
| M2-N3 | ACACCCAATGACGGCATA | 1.0000 | 0.0000 | 0.0000 | 0.9556 |
| M2-N4 | ACACACCACGTCCTAATG | 1.0000 | 0.0000 | 0.0000 | 0.83708 |
| M2-N5 | GCACAACACTCGACGCGG | 1.0000 | 0.0000 | 0.0000 | 0.93955 |
| M2-N6 | ACACACCGATCCCACTTG | 1.0000 | 0.0000 | 0.0000 | 1.09717 |
| M2-N7 | ACACGACACCACTTCCGA | 1.0000 | 0.0000 | 0.0000 | 0.91553 |
| M2-N8 | ACACACCGAAGCCGCCAT | 1.0000 | 0.0000 | 0.0000 | 1.30859 |
| M2-N9 | ACACACCGATCCCCATCG | 1.0000 | 0.0000 | 0.0000 | 0.60508 |
| M2-N10 | ACAACCCAACTCCGCTCG | 1.0000 | 0.0000 | 0.0000 | 0.99639 |
| M2-N11 | GCACAACACGGCAACCTT | 1.0000 | 0.0000 | 0.0000 | 1.05455 |
| M2-N12 | GCACACCCGATCCATCCG | 1.0000 | 0.0000 | 0.0000 | 0.97282 |
| M2-N13 | AGACGACACCTCACCGCT | 1.0000 | 0.0000 | 0.0000 | 0.50411 |
| M2-N14 | ACACCCTAACACTGCACG | 1.0000 | 0.0000 | 0.0000 | 1.02757 |
| M2-N15 | GAACCACACATCCGCATG | 1.0000 | 0.0000 | 0.0000 | 0.51354 |
| M2-N16 | ACATGCCACAACACCGAT | 1.0000 | 0.0000 | 0.0000 | 0.97921 |
| M2-N17 | ACACAACGCTGCTCTCCG | 1.0000 | 0.0000 | 0.0000 | 0.64343 |
| M2-N18 | ACAGGCCACCTCACCGAT | 1.0000 | 0.0000 | 0.0000 | 0.7973 |
| M2-N19 | ACACAGTCCGACCGTCTG | 1.0000 | 0.0000 | 0.0000 | 0.60328 |
| M2-N20 | ACACCCAAACTCCGCTCT | 1.0000 | 0.0000 | 0.0000 | 1.22111 |

| Seq ID | Sequence | Probability (class 0) | Probability (class 1) | Normalized probability | ΔF/F (1195 nm) |
|---|---|---|---|---|---|
| M2-P1 | AACCCTACACCATCCACA | 0.0020 | 0.9980 | 0.9980 | 0.98824 |
| M2-P2 | AACACAGCACCTTCCGAT | 0.0035 | 0.9965 | 0.9965 | 0.74183 |
| M2-P3 | AACCCGATCCAACCTCAT | 0.0041 | 0.9959 | 0.9959 | 1.0448 |
| M2-P4 | AACCAATCAACGTCCGCT | 0.0044 | 0.9956 | 0.9956 | 0.82122 |
| M2-P5 | AGCCCTTCACCACCAACT | 0.0084 | 0.9916 | 0.9916 | 2.91696 |
| M2-P6 | AACCACACCGCTCGTCCT | 0.0091 | 0.9909 | 0.9909 | 0.79546 |
| M2-P7 | AGCACCACCACATCCGAT | 0.0116 | 0.9884 | 0.9884 | 0.86068 |
| M2-P8 | AGCAACACGACTCCTGCT | 0.0129 | 0.9871 | 0.9871 | 1.85963 |
| M2-P9 | AACCGCATCCATTCCGCT | 0.0146 | 0.9854 | 0.9854 | 0.44503 |
| M2-P10 | TGCACGACACCTCTTCCT | 0.0152 | 0.9848 | 0.9848 | 1.03718 |
| M2-P11 | AACACAACTCGACGCGCA | 0.0176 | 0.9824 | 0.9824 | 1.57509 |
| M2-P12 | ACGAACACGGCACACACT | 0.0183 | 0.9817 | 0.9817 | 0.5442 |
| M2-P13 | ACGAACACTGCTCCTCCT | 0.0209 | 0.9791 | 0.9791 | 0.48006 |
| M2-P14 | AACCCGATCCATGCACCT | 0.0247 | 0.9753 | 0.9753 | 0.84969 |
| M2-P15 | AACACCAACCATTCCGCT | 0.0249 | 0.9751 | 0.9751 | 0.89558 |
| M2-P16 | AGCACCTGCACTCATCCT | 0.0250 | 0.9750 | 0.9750 | 0.78861 |
| M2-P17 | AACACAAGACAACGCGTG | 0.0292 | 0.9708 | 0.9708 | 2.53844 |
| M2-P18 | ACGCCAACCGATCCTCCG | 0.0304 | 0.9696 | 0.9696 | 0.76612 |
| M2-P19 | AGCACACCCGCTCCTACG | 0.0343 | 0.9657 | 0.9657 | 0.83824 |
| M2-P20 | ACGCCAACACGATGTGCT | 0.0374 | 0.9626 | 0.9626 | 0.76672 |

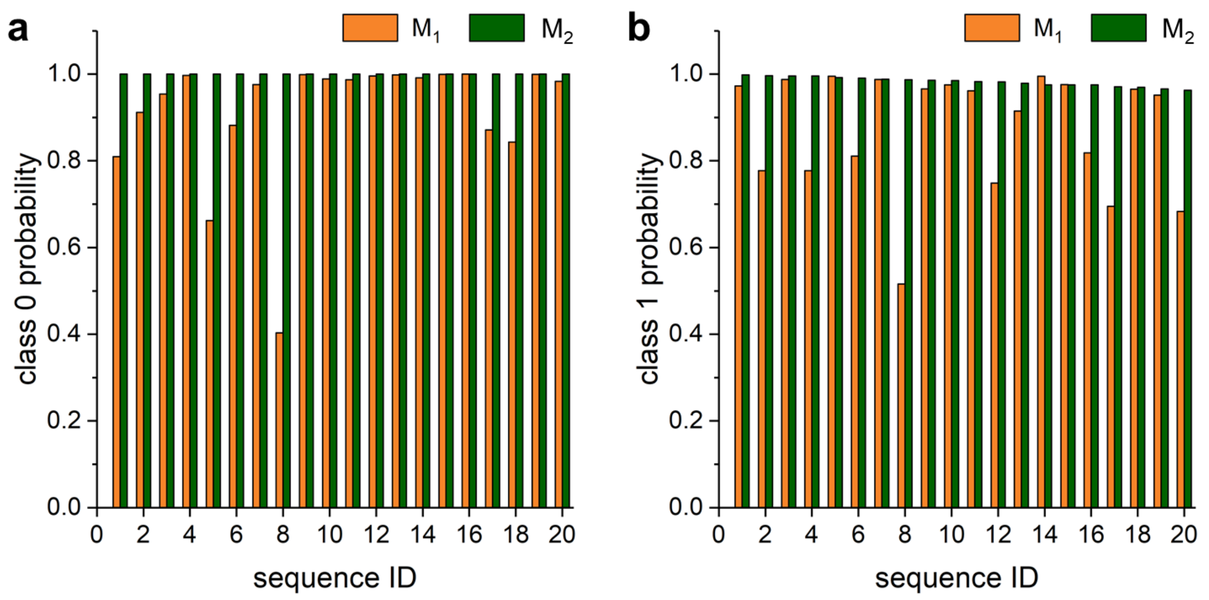


**Figure S4. Probabilities of sequences to be low or high responders to serotonin, as predicted by model $M_2$.** Class 0 or class 1 probabilities of 40 ssDNA sequences identified by model $M_2$ to be either low response-negative (a) or high response-positive (b) to serotonin. The probabilities obtained for those sequences by model $M_1$ are also shown for comparison.

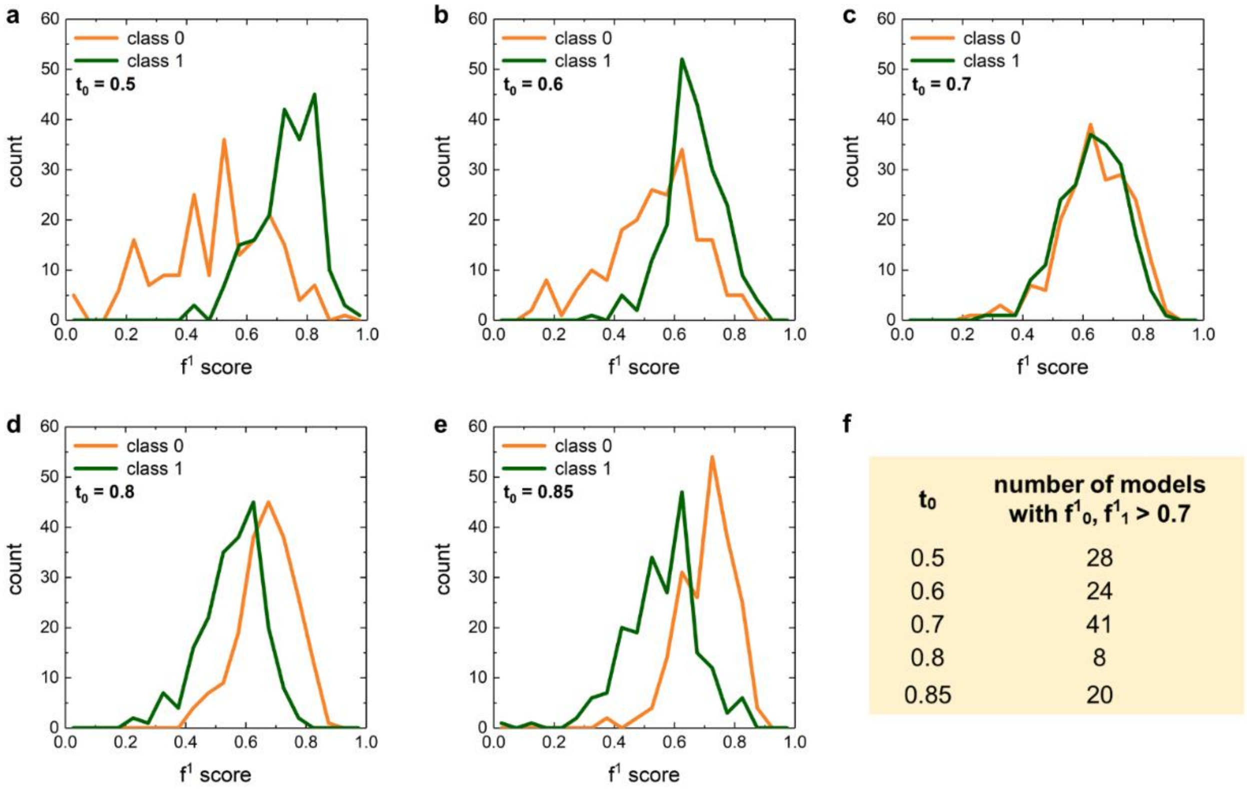


**Figure S5. Distribution of $f^1$ score values for ensembles of models trained and tested on expanded datasets.** Distribution of $f^1$ score values for 200 CNN models, obtained using different random states for different $t_0$ threshold values (a-e) for the expanded dataset. f) Number of models with $f^1$ scores > 0.7 when predicting class 0 and class 1 sequences.

**Table S6. Performance of CNN models with sequences in psv$_1$ format for several dataset classifications.** 200 models generated by using different random states (different training/testing dataset splits) are evaluated for performance and reliability. $t_0$ and $t_1$ are threshold values defining classes 0 and 1. Count is defined as the number of models (out of 200 models) for which TN, FP, FN, TP are all non-zero for the testing sequences. If either of those values are 0, some evaluation metrics automatically assume values of 0 or 1, which prevents model quality assessment. All the listed quantities are obtained from 200 models trained on data classified according to the stated threshold values.

| $t_1$ > 0.9, $t_0$ < 0.85, count = 196 | Ac | Prec$_0$ | Prec$_1$ | R$_0$ | R$_1$ | $f^1_0$ | $f^1_1$ | AUC$_0$ | AUC$_1$ | TN | FP | FN | TP |
|---|---|---|---|---|---|---|---|---|---|---|---|---|---|
| **mean** | 0.66 | 0.74 | 0.56 | 0.72 | 0.56 | 0.72 | 0.55 | 0.69 | 0.69 | 12.96 | 5.04 | 4.79 | 6.21 |
| **std. deviation** | 0.08 | 0.07 | 0.12 | 0.13 | 0.17 | 0.08 | 0.12 | 0.10 | 0.09 | 2.31 | 2.31 | 1.86 | 1.86 |
| **min** | 0.41 | 0.54 | 0.25 | 0.33 | 0.09 | 0.45 | 0.13 | 0.37 | 0.35 | 6 | 1 | 1 | 1 |
| **25% percentile** | 0.59 | 0.69 | 0.47 | 0.61 | 0.45 | 0.67 | 0.48 | 0.63 | 0.63 | 11 | 3 | 3 | 5 |
| **50% percentile** | 0.66 | 0.73 | 0.56 | 0.72 | 0.55 | 0.73 | 0.56 | 0.69 | 0.68 | 13 | 5 | 5 | 6 |
| **75% percentile** | 0.72 | 0.79 | 0.64 | 0.83 | 0.73 | 0.78 | 0.64 | 0.76 | 0.76 | 15 | 7 | 6 | 8 |
| **max** | 0.83 | 0.93 | 0.88 | 0.94 | 0.91 | 0.87 | 0.78 | 0.92 | 0.89 | 17 | 12 | 10 | 10 |

| $t_1 > 0.9$, $t_0 < 0.8$, count = 196 | Ac | $Prec_0$ | $Prec_1$ | $R_0$ | $R_1$ | $f^1_0$ | $f^1_1$ | $AUC_0$ | $AUC_1$ | TN | FP | FN | TP |
|---|---|---|---|---|---|---|---|---|---|---|---|---|---|
| **mean** | 0.64 | 0.72 | 0.55 | 0.68 | 0.58 | 0.69 | 0.55 | 0.67 | 0.67 | 11.5 | 5.47 | 4.66 | 6.34 |
| **standard deviation** | 0.08 | 0.08 | 0.11 | 0.13 | 0.15 | 0.09 | 0.11 | 0.10 | 0.10 | 2.26 | 2.26 | 1.69 | 1.69 |
| **min** | 0.29 | 0.41 | 0.09 | 0.35 | 0.09 | 0.41 | 0.09 | 0.35 | 0.32 | 6 | 1 | 1 | 1 |
| **25% percentile** | 0.61 | 0.67 | 0.50 | 0.59 | 0.45 | 0.64 | 0.50 | 0.62 | 0.61 | 10 | 4 | 3.75 | 5 |
| **50% percentile** | 0.64 | 0.71 | 0.55 | 0.71 | 0.55 | 0.71 | 0.56 | 0.68 | 0.68 | 12 | 5 | 5 | 6 |
| **75% percentile** | 0.68 | 0.75 | 0.60 | 0.76 | 0.66 | 0.74 | 0.63 | 0.74 | 0.73 | 13 | 7 | 6 | 7.25 |
| **max** | 0.82 | 0.90 | 0.83 | 0.94 | 0.91 | 0.86 | 0.76 | 0.93 | 0.89 | 16 | 11 | 10 | 10 |

| $t_1 > 0.9$, $t_0 < 0.7$, count = 192 | Ac | $Prec_0$ | $Prec_1$ | $R_0$ | $R_1$ | $f^1_0$ | $f^1_1$ | $AUC_0$ | $AUC_1$ | TN | FP | FN | TP |
|---|---|---|---|---|---|---|---|---|---|---|---|---|---|
| **mean** | 0.64 | 0.69 | 0.61 | 0.64 | 0.64 | 0.66 | 0.62 | 0.69 | 0.68 | 8.36 | 4.64 | 3.92 | 7.08 |
| **standard deviation** | 0.09 | 0.10 | 0.10 | 0.15 | 0.16 | 0.10 | 0.11 | 0.10 | 0.10 | 1.9 | 1.9 | 1.79 | 1.79 |
| **min** | 0.42 | 0.46 | 0.36 | 0.23 | 0.18 | 0.32 | 0.25 | 0.43 | 0.41 | 3 | 1 | 1 | 2 |
| **25% percentile** | 0.58 | 0.62 | 0.55 | 0.54 | 0.55 | 0.59 | 0.55 | 0.62 | 0.63 | 7 | 3 | 3 | 6 |
| **50% percentile** | 0.63 | 0.69 | 0.60 | 0.69 | 0.64 | 0.67 | 0.64 | 0.69 | 0.69 | 9 | 4 | 4 | 7 |
| **75% percentile** | 0.71 | 0.75 | 0.69 | 0.77 | 0.73 | 0.73 | 0.69 | 0.76 | 0.76 | 10 | 6 | 5 | 8 |
| **max** | 0.83 | 0.91 | 0.89 | 0.92 | 0.91 | 0.86 | 0.83 | 0.91 | 0.91 | 12 | 10 | 9 | 10 |

| $t_1 > 0.9$, $t_0 < 0.6$, count = 191 | Ac | $Prec_0$ | $Prec_1$ | $R_0$ | $R_1$ | $f^1_0$ | $f^1_1$ | $AUC_0$ | $AUC_1$ | TN | FP | FN | TP |
|---|---|---|---|---|---|---|---|---|---|---|---|---|---|
| **mean** | 0.62 | 0.59 | 0.65 | 0.53 | 0.69 | 0.54 | 0.66 | 0.66 | 0.65 | 4.75 | 4.25 | 3.42 | 7.58 |
| **standard deviation** | 0.09 | 0.12 | 0.09 | 0.18 | 0.16 | 0.14 | 0.10 | 0.12 | 0.11 | 1.62 | 1.62 | 1.76 | 1.76 |
| **min** | 0.35 | 0.25 | 0.33 | 0.11 | 0.18 | 0.15 | 0.24 | 0.32 | 0.35 | 1 | 1 | 1 | 2 |
| **25% percentile** | 0.55 | 0.50 | 0.59 | 0.44 | 0.64 | 0.46 | 0.60 | 0.59 | 0.59 | 4 | 3 | 2 | 7 |
| **50% percentile** | 0.60 | 0.57 | 0.64 | 0.56 | 0.73 | 0.56 | 0.67 | 0.67 | 0.65 | 5 | 4 | 3 | 8 |
| **75% percentile** | 0.70 | 0.67 | 0.71 | 0.67 | 0.82 | 0.63 | 0.73 | 0.74 | 0.73 | 6 | 5 | 4 | 9 |
| **max** | 0.90 | 0.89 | 0.91 | 0.89 | 0.91 | 0.89 | 0.91 | 0.99 | 0.94 | 8 | 8 | 9 | 10 |

| $t_1 > 0.9$, $t_0 < 0.5$, count = 180 | Ac | $Prec_0$ | $Prec_1$ | $R_0$ | $R_1$ | $f^1_0$ | $f^1_1$ | $AUC_0$ | $AUC_1$ | TN | FP | FN | TP |
|---|---|---|---|---|---|---|---|---|---|---|---|---|---|
| **mean** | 0.64 | 0.55 | 0.70 | 0.50 | 0.73 | 0.51 | 0.71 | 0.67 | 0.67 | 3.51 | 3.49 | 2.95 | 8.05 |
| **standard deviation** | 0.09 | 0.14 | 0.09 | 0.21 | 0.15 | 0.16 | 0.10 | 0.12 | 0.13 | 1.44 | 1.44 | 1.63 | 1.63 |
| **min** | 0.44 | 0.25 | 0.55 | 0.14 | 0.18 | 0.18 | 0.29 | 0.30 | 0.29 | 1 | 1 | 1 | 2 |
| **25% percentile** | 0.61 | 0.50 | 0.64 | 0.29 | 0.64 | 0.40 | 0.67 | 0.60 | 0.60 | 2 | 2 | 2 | 7 |
| **50% percentile** | 0.67 | 0.56 | 0.69 | 0.50 | 0.73 | 0.51 | 0.73 | 0.68 | 0.69 | 3.5 | 3.5 | 3 | 8 |
| **75% percentile** | 0.72 | 0.67 | 0.77 | 0.71 | 0.82 | 0.63 | 0.77 | 0.77 | 0.77 | 5 | 5 | 4 | 9 |
| **max** | 0.89 | 0.86 | 0.91 | 0.86 | 0.91 | 0.86 | 0.91 | 0.94 | 0.95 | 6 | 6 | 9 | 10 |

**Table S7.** DNA sequences predicted by combined models to belong to either class 0 or class 1, and their experimental ΔF/F response to serotonin. The sequences predicted to be in class 1 are highlighted in green. The sequences that are experimentally validated to belong to class 0 (top table) or 1 (bottom table) are highlighted in orange and blue, respectively. All the sequences are flanked by two $C_6$-mer PCR primers on each side.

| Seq ID | Sequence | Probability (class 0) | Probability (class 1) | Predicted ΔF/F | ΔF/F (exp, 1195 nm) |
|---|---|---|---|---|---|
| E3-N1 | ACACCACAGCACTCCGAT | 0.9407 | 0.0602 | 0.4801 | 0.70365 |
| E3-N2 | ACACCCAACGTCTGCTCT | 0.9296 | 0.0604 | 0.6816 | 0.46680 |
| E3-N3 | ACACCCTAACTCCGCTCT | 0.9355 | 0.0671 | 0.6393 | 1.27976 |
| E3-N4 | ACACCCTGAGTCCGCACA | 0.9509 | 0.0469 | 0.6569 | 0.51381 |
| E3-N5 | ACACACCGATCCACCGCT | 0.9634 | 0.0368 | 0.4362 | 0.80368 |
| E3-N6 | ACATACCCACTCCGCTCG | 0.9651 | 0.0372 | 0.4898 | 0.58693 |
| E3-N7 | GCACACCGATCCTACCAG | 0.9754 | 0.0220 | 0.4543 | 0.60202 |
| E3-N8 | ACACACCCTAATCTCGCT | 0.9324 | 0.0577 | 0.5526 | 0.56565 |
| E3-N9 | ACAAACCGCTCATCCGAT | 0.9373 | 0.0617 | 0.4783 | 0.46788 |
| E3-N10 | ACACTCCGACCCTTCTCG | 0.9405 | 0.0683 | 0.4963 | 0.70365 |
| E3-P1 | AACGCCACCCTAACTCCG | 0.0198 | 0.9823 | 1.3064 | 1.57806 |
| E3-P2 | AGCCCGAACCAGACACCG | 0.0198 | 0.9736 | 1.3454 | 0.56539 |
| E3-P3 | AACCCGAACCTAACTGCG | 0.0191 | 0.9735 | 1.3291 | 0.43275 |
| E3-P4 | AACGCAACACGACCTGTG | 0.0294 | 0.9675 | 1.3142 | 0.46865 |
| E3-P5 | AACCCAGACCGACCACCT | 0.0317 | 0.9633 | 1.3936 | 1.41762 |
| E3-P6 | GACCCAAAGCCAACACCT | 0.0471 | 0.9528 | 1.3934 | 2.07183 |
| E3-P7 | GACCCTAACACAGCACCA | 0.0158 | 0.9867 | 1.3615 | 1.30689 |
| E3-P8 | AACCCTAACCGATCACTG | 0.0208 | 0.9767 | 1.3633 | 1.32151 |
| E3-P9 | AACACGACCCGACCTGTG | 0.0142 | 0.9740 | 1.3852 | 0.53283 |
| E3-P10 | GACCCAAACCTACCTCCA | 0.0339 | 0.9655 | 1.4481 | 1.57806 |

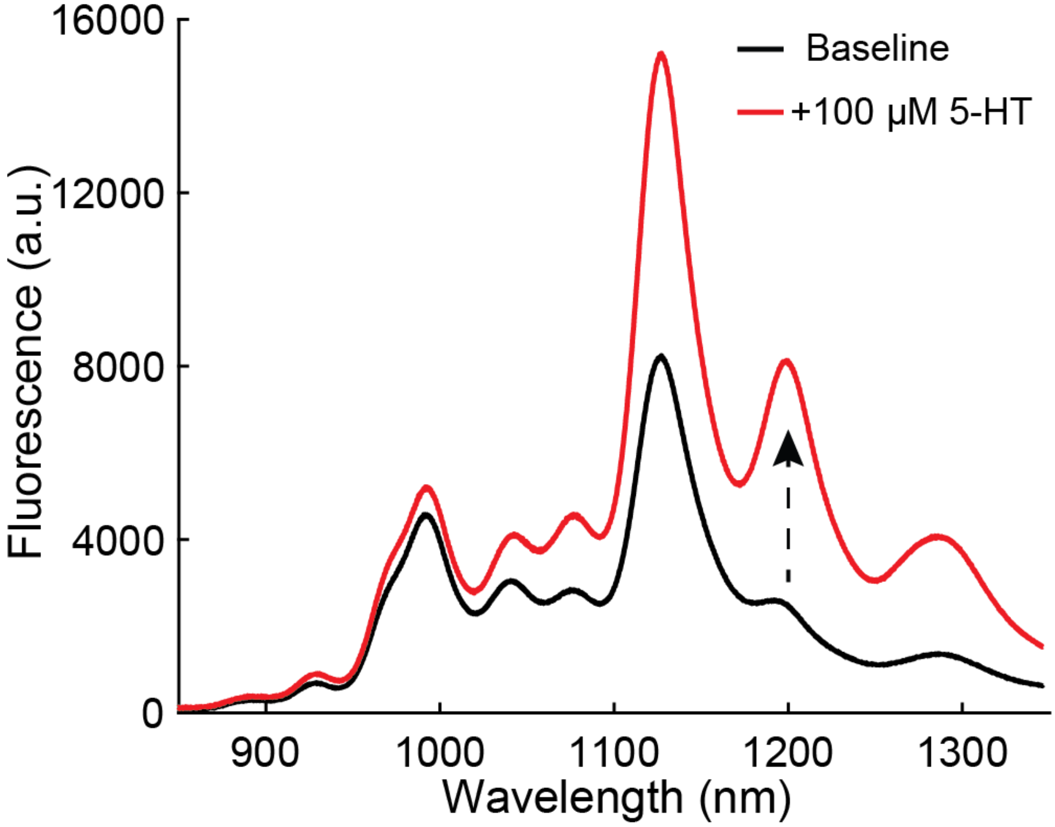


**Figure S6. Example spectra of one of the newly identified DNA-SWNT sensors.** Fluorescence spectra of E3-P6 sequence before the addition of serotonin (5-HT) to DNA-SWNT suspension (black trace) and after the addition of 100 µM serotonin (red trace).

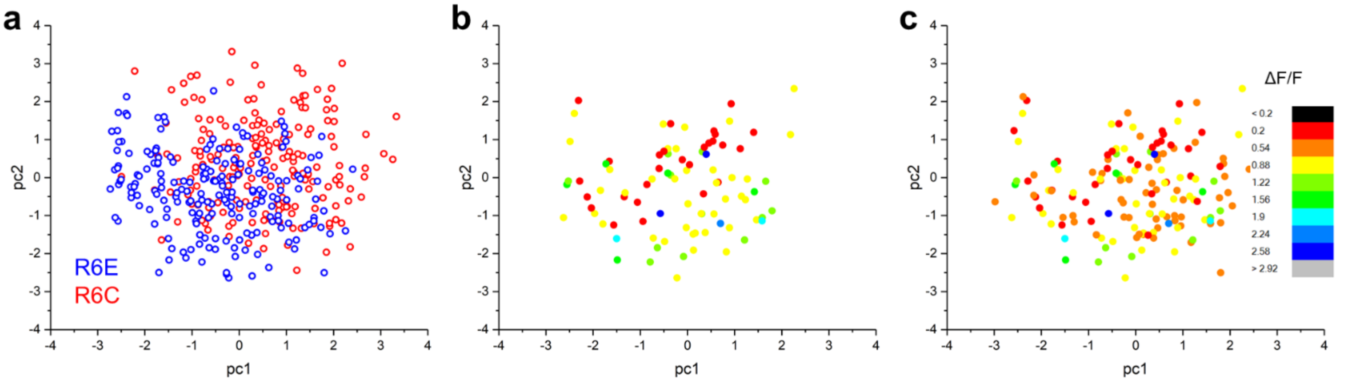


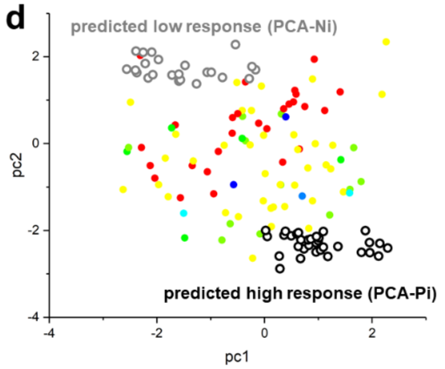


**e**

| | | |
|---|---|---|
| PCA-N1 | TACCACTCCAACTCCGCT | 0.9112 |
| PCA-N2 | ACACGCCCTAACTCCGCT | 1.3866 |
| PCA-N3 | AGGCACACTCATCCGCGT | 0.7582 |
| PCA-N4 | ACACCGCCTAATGCACCT | 1.1045 |
| PCA-N5 | CACGATCCTACTCACGCT | 0.9905 |
| PCA-N6 | ACACCCGATCCTGCACCG | 1.5967 |
| PCA-N7 | AACCAACATCCTTCCGCT | 0.5780 |
| PCA-N8 | GGCAACGATACTCAACCT | 0.9290 |
| PCA-N9 | ACCACACCTCATTCCGCT | 0.7434 |
| PCA-N10 | AACACCGATCCTCCATGT | 0.9112 |

| | | |
|---|---|---|
| PCA-P1 | ACACACAGACTCACCGCT | 1.3136 |
| PCA-P2 | ACACCGATCCTACGCACT | 0.6375 |
| PCA-P3 | GCCACCAGACTCAATGCT | 0.8922 |
| PCA-P4 | ACACCGATCCTAACTCCG | 0.6160 |
| PCA-P5 | AACACAACATTCCGCTCG | 0.5315 |
| PCA-P6 | AACCTCACACTCATTCCG | 0.4846 |
| PCA-P7 | ACCACGATCCTCACTCCG | 0.3629 |
| PCA-P8 | ACGACCCTATGAACCTCG | 0.6053 |
| PCA-P9 | GACCACAGAATACGGCCT | 0.8912 |
| PCA-P10 | AGCAACAGACGACCTGCT | 1.4129 |

**Figure S7. Principal component analyses of sequences.** a) Principal component analyses for the top 200 sequences from the R6E and R6C datasets. b) Sequences from the expanded dataset with ΔF/F>0.9 and ΔF/F<0.5, colored according to their ΔF/F values. The sequences are plotted in the PC space determined in the analysis from panel a. c) Sequences from the expanded dataset with ΔF/F>0.9 and ΔF/F<0.85, colored according to their ΔF/F values. The legend is the same as in panel c. The sequences are plotted in the PC space determined in the analysis from panel a. d) Same plot as in panel b, with 20 sequences predicted to have low response and 20 sequences predicted to have high response, according to the regions in the plot. e) The optical response ΔF/F of ssDNA-SWNT conjugates to 100 µM serotonin, obtained for 20 ssDNA sequences selected from the grey and black datapoints in panel d.

**Table S8. Analyses of motifs in DNA sequences.** Base motifs in DNA sequences with ΔF/F > 0.9 and their frequencies from the expanded dataset used to train and test model $M_2$, identified when maximal occurrence frequency of negative sequences, $f_N$, is set to zero. The motifs were found by using the MERCI software[2].

| Motif | Frequency |
|---|---|
| AA_A | 3 |
| AACAC_GC | 3 |
| AACA_C | 4 |
| AACA_T | 3 |
| AAC_A | 4 |
| AACCA | 3 |
| AACCA_A | 3 |
| AACCC | 3 |
| AACCG | 3 |
| AACC_G | 5 |
| AACC_GA | 3 |
| AAC_CC | 4 |
| AAC_CG | 4 |
| AA_CA | 4 |
| AA_CCC | 3 |
| AA_CCG | 4 |
| A_ACCAA | 3 |
| ACAACA | 3 |
| ACAAC_C | 4 |
| ACAC_ATT | 3 |
| ACAC_ATTC | 3 |
| ACAC_ATTCC | 3 |
| ACAC_ATTCCG | 3 |
| ACAC_ATTCCGC | 3 |
| ACACCAA | 3 |
| ACACG_C | 3 |
| ACAC_TT | 3 |
| ACAC_TTC | 3 |
| ACAC_TTCC | 3 |
| ACAC_TTCCG | 3 |
| ACAC_TTCCGC | 3 |
| ACA_CCAA | 3 |
| ACAG_AC | 3 |
| ACA_TG | 3 |
| ACA_TGT | 3 |
| AC_ACCAA | 3 |
| AC_AGAC | 3 |
| ACCA_TC | 3 |

| Motif | Frequency |
|---|---|
| ACC_TC | 3 |
| ACC_TCC | 3 |
| AC_CAAC | 3 |
| AC_CATT | 3 |
| AC_CATTC | 3 |
| AC_CATTCC | 3 |
| AC_CATTCCG | 3 |
| AC_CATTCCGC | 3 |
| ACGACA | 3 |
| ACGACA_A | 3 |
| ACGC_C | 3 |
| AC_GA | 3 |
| AC_GTCC | 3 |
| ACTCCA | 3 |
| ACT | 3 |
| A_CGTC | 3 |
| A_CGTCC | 3 |
| AGACG | 3 |
| AGAC_TC | 3 |
| AG_ACA | 3 |
| AG_ACAA | 3 |
| AGCACA_C | 3 |
| AGCACA_CA | 3 |
| AGCACA_CAC | 3 |
| AGCA | 3 |
| AGG | 3 |
| A_GCACA | 4 |
| A_GCACAA | 3 |
| A_GG | 3 |
| ATCCA | 3 |
| A_TCCA | 3 |
| CAACAC | 4 |
| CAACA_C | 4 |
| CAACA | 3 |
| CAAC_A | 3 |
| CAACC | 3 |
| CAACC | 5 |
| CAACC_G | 3 |

| Motif | Frequency |
|---|---|
| CAAC_C | 6 |
| CAA_CA | 3 |
| CA_ACCAA | 3 |
| CACAACA | 3 |
| CACAAC_C | 4 |
| CACA_CAC | 3 |
| CACAG_AC | 3 |
| CACCAA | 3 |
| CACC_TC | 3 |
| CACC_TCC | 3 |
| CACG_C | 3 |
| CA_CCAA | 3 |
| CAGACG | 3 |
| CAGAC_TC | 3 |
| CAT_CA | 3 |
| CAT_CAC | 3 |
| C_ACCAA | 3 |
| CCAACAC | 3 |
| CCACACC | 3 |
| CC_CACCA | 3 |
| C_CAACC | 3 |
| C_CAGAC | 3 |
| C_CGGC | 3 |
| CGA_AC | 3 |
| CGA_ACA | 3 |
| CGACA | 3 |
| CGACA_A | 3 |
| CG_A | 3 |
| CGCA_ | 3 |
| C_GA | 4 |
| C_GCGT | 3 |
| CTCCA | 3 |
| GA_AC | 3 |
| GA_ACA | 3 |
| GACA_A | 4 |
| GAC_C | 6 |
| GAC_CA | 5 |
| G_ACC | 3 |

| Motif | Frequency |
|---|---|
| G_ACT | 3 |
| GCACAAC_C | 3 |
| GCACA_CAC | 3 |
| GCA_CC | 3 |
| GC_ACC | 3 |
| GC_CAACA | 3 |
| GC_CC | 3 |
| G_CAAC | 4 |
| GGA_C | 3 |
| GG_AC | 3 |
| GGCA_C | 3 |
| GG_CAAC | 3 |
| TC_A | 4 |
| TC_AA | 3 |
| TCCA | 3 |
| TCC_A | 3 |
| TC_CA | 3 |
| T_CAA | 3 |
| T_CCA | 3 |
